# Coordinated overexpression of *OsSUT1, OsSWEET11* and *OsSWEET14* in rice impairs carbohydrate metabolism that has implications in plant growth, yield and susceptibility to *Xanthomonas oryzae pv oryzae (Xoo)*

**DOI:** 10.1101/2021.01.07.425507

**Authors:** Jitender Singh, Donald James, V Mohan Murali Achary, Manish Kumar Patel, Jitendra Kumar Thakur, M K Reddy, Baishnab Charan Tripathy

## Abstract

Enhancing carbohydrate export to sink tissues is considered as a feasible approach for improving photosynthetic efficiency and crop yield. In *Oryza sativa* Sucrose Transporter OsSUT1 located in companion cells and Sugars Will Eventually be Exported Transporters (SWEETs); OsSWEET11 and OsSWEET14 present in phloem parenchyma mesophyll cell plasma membranes are involved in long distance sucrose transport. *OsSWEET11* and *OsSWEET14* also play important role in host-pathogen interaction of rice plants and *Xanthomonas oryzae* pv *oryzae* (*Xoo*) that causes bacterial leaf blight. Three genes, *OsSUT1, OsSWEET11*, and *OsSWEET14* were overexpressed under the control of their native promoters in rice to modulate long distance sugar transport and disease resistance. The transgenics displayed several phenotypic aberrations such as reduced plant height and seed weight due to altered sucrose transport and metabolism. Lower sucrose transport rate in transgenics than the WT resulted in reduced sucrose, fructose and glucose and increased starch accumulation in their leaves at the end of dark period. Transcriptional analysis revealed a reduction in the expression of genes involved in sucrose synthesis pathway in transgenics. Normal growth and development of transgenic seedlings were restored in growth media supplemented with 3% sucrose demonstrating *in planta* sucrose limitation. Remarkably, transgenic lines had diminished susceptibility to *Xoo* than the WTs due to low sugar content in the leaves demonstrating that rice plants maintain an optimum level of SWEETs for proper plant growth and development, and upregulation of these *SWEETs* in rice mimicks *Xoo* attack impelling plants to reduce sugar content in the apoplasm to inhibit pathogen growth.

## Introduction

SWEETs are transmembrane proteins responsible for sugar transport, copper transport, nectar secretion, pollen development, seed filling, senescence, host-pathogen interactions and biotic and abiotic stress tolerance in plants (Chandran, 2015; Sosso et al., 2015; Miao et al., 2017; Jeena et al., 2019). In higher plants SWEETs are divided into four clades (I, II, III, and IV), and SWEETs belonging to clade III function as sucrose transporters (Chen et al., 2012; Jeena et al., 2019). Long distance sucrose transport from source (leaves) to sink tissues such as seeds, flowers, stems and roots is essential for proper plant growth and development. In rice, *Arabidopsis* and several other plants, first step in long distance sucrose translocation is the SWEET proteins mediated efflux of sucrose from the mesophyll cells and vascular parenchyma to the apoplasm followed by sucrose influx to phloem cells against the concentration gradient across the plasma membrane of cells making up the sieve element/companion cell complex by sucrose/H^+^ symporter (SUT1) (Chen et al., 2012). Both OsSWEET11 and OsSWEET14 of clade III function as low affinity glucose and sucrose transporters and are involved in efflux of sucrose from the parenchyma cells to the apoplasm in rice, thus ascertaining their role in transport of sucrose over long distances from leaves to regions of growth and storage such as roots and seeds (Chen et al., 2010; Yuan and Wang, 2013). Rice T-DNA insertional mutants of *Ossweet14* displayed dwarf phenotype and had smaller seeds compared to the WTs (Antony et al., 2010; Wu et al., 2018), indicating impaired sucrose transport from leaves to seeds. Disruption in the coding sequence and the EBEs (effector binding elements) in the promoter of *OsSWEET14* by genome editing technologies, however, did not reveal any recognizable phenotypic difference as compared to the WTs (Li et al., 2012; Blanvillain-Baufumé et al., 2017; Eom et al., 2019; Oliva et al., 2019; Xu et al., 2019; Kim et al., 2020; Zeng et al., 2020). In fact, height of the knock out mutants of *Ossweet14* was slightly higher than the WTs (Zeng et al., 2020). Knock out mutants of *Ossweet11* are defective in grain filling and the seeds show delayed germination and growth as compared to WTs (Ma et al., 2017; Yang et al., 2018). Pollens of *Ossweet11* RNAi rice plants were found defective and had reduced viability (Yang et al., 2006). Similarly, *Atsweet11;12* (*AtSWEET11* and *AtSWEET12* are orthologs of *OsSWEET11* and *OsSWEET14*) *Arabidopsis* double mutant plants were smaller in size and had reduced seed weight due to their weakened ability to export sucrose from leaves to sink organs (Chen et al., 2015).

*Xoo* is the causal agent of bacterial rice leaf blight. *Xoo* induces transcription of both *OsSWEET11* and *OsSWEET14* to increase the availability of sugar in vascular tissues for its survival and growth (Zaka et al., 2018). OsSWEET13 is another member of clade III, which is exploited by *Xoo* for its virulence (Zaka et al., 2018; Oliva et al., 2019; Xu et al., 2019).

Rice genome encodes 5 SUTs; *OsSUT1, OsSUT2, OsSUT3, OsSUT4*, and *OsSUT5* (Aoki et al., 2003). OsSUT2 is localized in tonoplast membrane, while all other are plasma membrane proteins (Eom et al., 2011). OsSUT3, OsSUT4, and OsSUT5 are preferentially expressed in sink leaves, whereas, OsSUT3 and OsSUT5 are minimally expressed in germinating seeds (Aoki et al., 2003; Sun et al., 2010). OsSUT3 and 4 are strongly expressed in pollens (Eom et al., 2012; Chung et al., 2014). Sequence and functional similarity analyses, and multiple experimental evidences suggest that OsSUT1 is the primary phloem loading SUT in rice. OsSUT1 is the ortholog of AtSUC2 and ZmSUT1, which are the key phloem loading proteins in *Arabidopsis* and maize respectively (Slewinski et al., 2009; Eom et al., 2016). Transcript expression profile and *proOsSUT1::GUS* localization studies reveal the presence of OsSUT1 in all the tissues involved in long distance sucrose transport *i*.*e*. flag leaf blade, sheath, internode, pollen and panicles (Aoki et al., 2003; Scofield et al., 2007a; Hirose et al., 2010). Out of the five rice *SUTs*, only *OsSUT1* under the regulation of *AtSUC2* promoter was able to complement the *Atsuc2* mutant phenotype of severe growth retardation, a typical feature of impaired phloem loading (Eom et al., 2016). In addition, *Ossut1* knock down rice plants showed impaired pollen function, grain filling and germination (Scofield et al., 2002; Scofield et al., 2007b; Hirose et al., 2010).

All three proteins *i*.*e*. OsSUT1, OsSWEET11, and OsSWEET14 are involved in phloem loading of sucrose in leaves, its long-distance transport to various sink organs and grain filling. Very recently, it has been found that the transcription factor OsDOF11 regulates the expression of rice *SWEET* and *SUT* genes (Wu et al., 2018). OsDOF11 directly binds to the promoters of *OsSUT1, OsSWEET11* and *OsSWEET14*, suggesting that these transporters work in conjunction to translocate sucrose from source to sink tissues (Wu et al., 2018). Mutants of *Osdof11* showed pleiotropic defects such as growth retardation, reduced tiller numbers and grains per panicle as compared to WT plants (Wu et al., 2018; Kim et al., 2020).

In this study, we attempted to modulate the long distance sucrose transportation and grain filling of rice plants by increasing the expression of *OsSUT1, OsSWEET11*, and *OsSWEET14* in a tissue specific manner by using their native promoters as proposed by us earlier (Singh et al., 2014).

## Results

### Cloning and in vitro pyramiding of *OsSUT1, OsSWEET11*, and *OsSWEET14* genes in plant transformation vector, its transformation in rice calli, generation of transgenic plants and their molecular confirmation

The nucleotide fragments of ∼3750 and ∼3690 bp for *OsSWEET11* and *14* genes respectively were PCR amplified from the genomic DNA (gDNA) of rice plants. The fragments comprise exons, introns, and the 1.5 kb region upstream of the ATG translation start site for promoter activity (Figure S1). The amplified products were successfully cloned in gateway compatible entry vectors (EVs) for further sub-cloning into plant transformation vector *pMDC99*. Full length coding sequence of *OsSUT1* gene was amplified by using cDNA of rice as template (Figure S1). Nucleotide fragment of 1600 bp upstream of the start codon ATG of the *OsSUT1* was amplified and fused to the 5’ end of the *OsSUT1* coding sequence to be used as promoter. The *CaMV 35S* terminator sequence was fused to the 3’ side of the coding sequences of all the three transgenes in EVs to function as the terminator signal for transcription.

All the three expression cassettes namely; *OsSWEET11* (in EV-1), *OsSWEET14* (in EV-2) and *OsSUT1* (in EV-1) were stacked together in *pMDC99* using the Multi-Round Gateway cloning technology (Figure 1A, Figure S1) (James et al., 2018). Over 100 putative T_0_ transgenic lines were produced. Most of the transformants were PCR positive and showed integration of all the transgenes (Figure S2). Southern analysis of 9 PCR positive T_2_ transgenic lines with *HPTII* as a probe was performed. Single to multiple hybridization bands with a distinct pattern for each transgenic line was observed suggesting independent integration of the transgenes in each transgenic event (Figure S2). No hybridization signal appeared in the WT. Three independent T_2_ transgenic lines (SW5, SW8, and SW9), which were PCR positive for all the transgenes and had single copy insertion of the recombinant construct, were selected for reverse transcriptase-PCR (RT-PCR) analysis. Amplification was observed in all the three transgenic lines indicating the expression of *OsSUT1, OsSWEET11* and *OsSWEET14* transgenes (Figure S2). The transcript abundance of all the transgenes was higher in SW5 and SW8 as compared to SW9. This was further confirmed by real time-quantitative PCR (RT-qPCR) analysis, which showed that the expression of *OsSUT1, OsSWEET11* and *OsSWEET14* is higher in SW5 as compared to SW9 (Figure 1B). In comparison to the WT, *OsSWEET11* was upregulated by over 8 and 11 fold in SW9 and SW5 respectively. Similarly, transcription of *OsSWEET14* was increased by a factor of 11 and 15 in SW9 and SW5, respectively (Figure 1B). Expression of *OsSUT1* was increased by more than 70 folds in SW9 and SW5 in comparison to the WT (Figure 1B). This could be due to the use of only coding sequence of *OsSUT1* in the recombinant construct as compared to the complete gene sequences of *OsSWEET11* and *OsSWEET14* that include both exons and introns.

**Figure 1.**
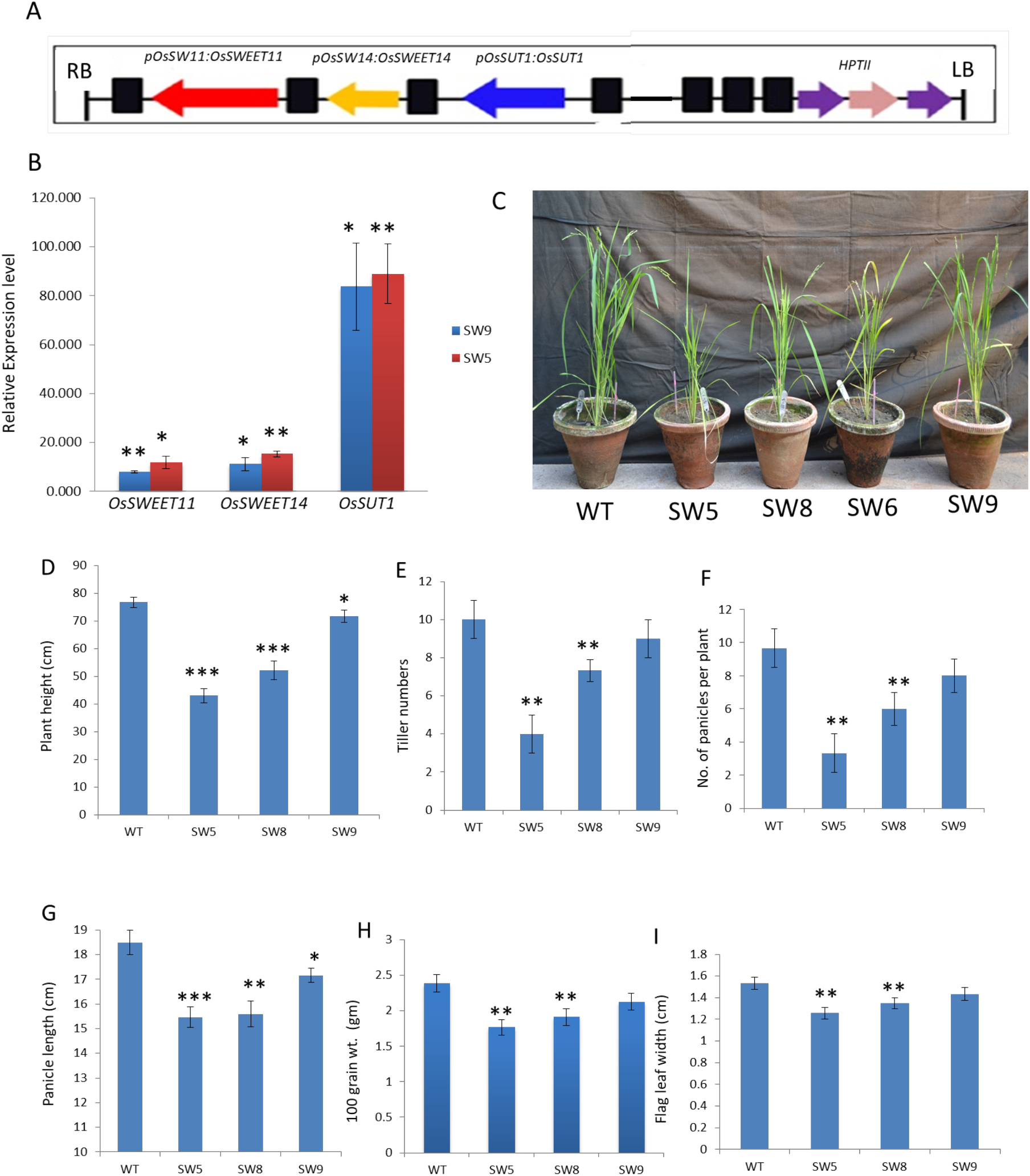
Phenotypic data analysis of WT and transgenic lines SW5, SW8, and SW9. **(A)** Final recombinant cassette in *pMDC99* transformed into the WT rice calli; **(B)** RT-qPCR analysis of *OsSWEET11, OsSWEET14* and *OsSUT1* in SW9 and SW5 lines **(C)** WT and different transgenic lines grown in green house; **(D-I)** Measurement of different morphological and agronomic features of WT and SW5, 8, and 9; **(D)** Plant height; **(E)** Tiller number; **(F)** Panicle number; **(G)** Panicle length; **(H)** Hundred grain weight; **(I)** Flag leaf width. Error bars show ± SEM. *P ≤ 0.05, **P ≤ 0.01, ***P ≤ 0.001

### Simultaneous enhancement in tissue specific expression of *OsSUT1, OsSWEET11*, and *OsSWEET14* leads to pleiotropic effects

All the transgenic plants harboring the recombinant cassette containing the three genes showed growth retardation (Figure 1C). SW9 showed mild whereas SW5 and SW8 showed very severe phenotypes as compared to the WT. SW5 and SW8 displayed semi-dwarf phenotype as compared to WT (Figure 1C). All the transformed lines had decreased plant height, tiller numbers, number of panicles per plant, panicle size, grain weight and flag leaf width than the WT (Figure 1D-I).

Transgenic lines SW5 and SW8 that displayed severe phenotypes (Figure 1C-I) had higher expression of *OsSWEET11* and *OsSWEET14* than SW9 that had a small impact on plant height and other growth parameters (Figure S2). RT-qPCR analysis also showed that the levels of *OsSWEET11* and *OsSWEET14* transcripts are higher in SW5 than SW9 (Figure 1B). This indicates that the intensity of the phenotypic aberrations is directly proportional to the level of expression of *OsSWEET11* and *14*.

### Sucrose transport is impaired in transgenic lines

The photosynthetic carbon assimilation rate measured in ambient CO_2_ was substantially reduced in SW5 (−40%) and SW9 (−15%) compared to the WT (Figure 2A). Decreased photosynthetic rates of these transgenic lines resulted in lower starch content in flag leaves of SW5 and SW9 than the WT, measured at the end of day (Figure 2B). However, starch content measured at the end of night period was significantly higher (∼200%) in both the transgenic lines than the WT (Figure 2B). Consequently, the transgenic lines had higher iodine staining of flag leaves harvested after end of the night (Figure 2C) indicating that sucrose transport in leaves of transgenic lines is reduced and the photosynthate produced during day is deposited in the form of starch in chloroplasts of the overexpressors.

**Figure 2.**
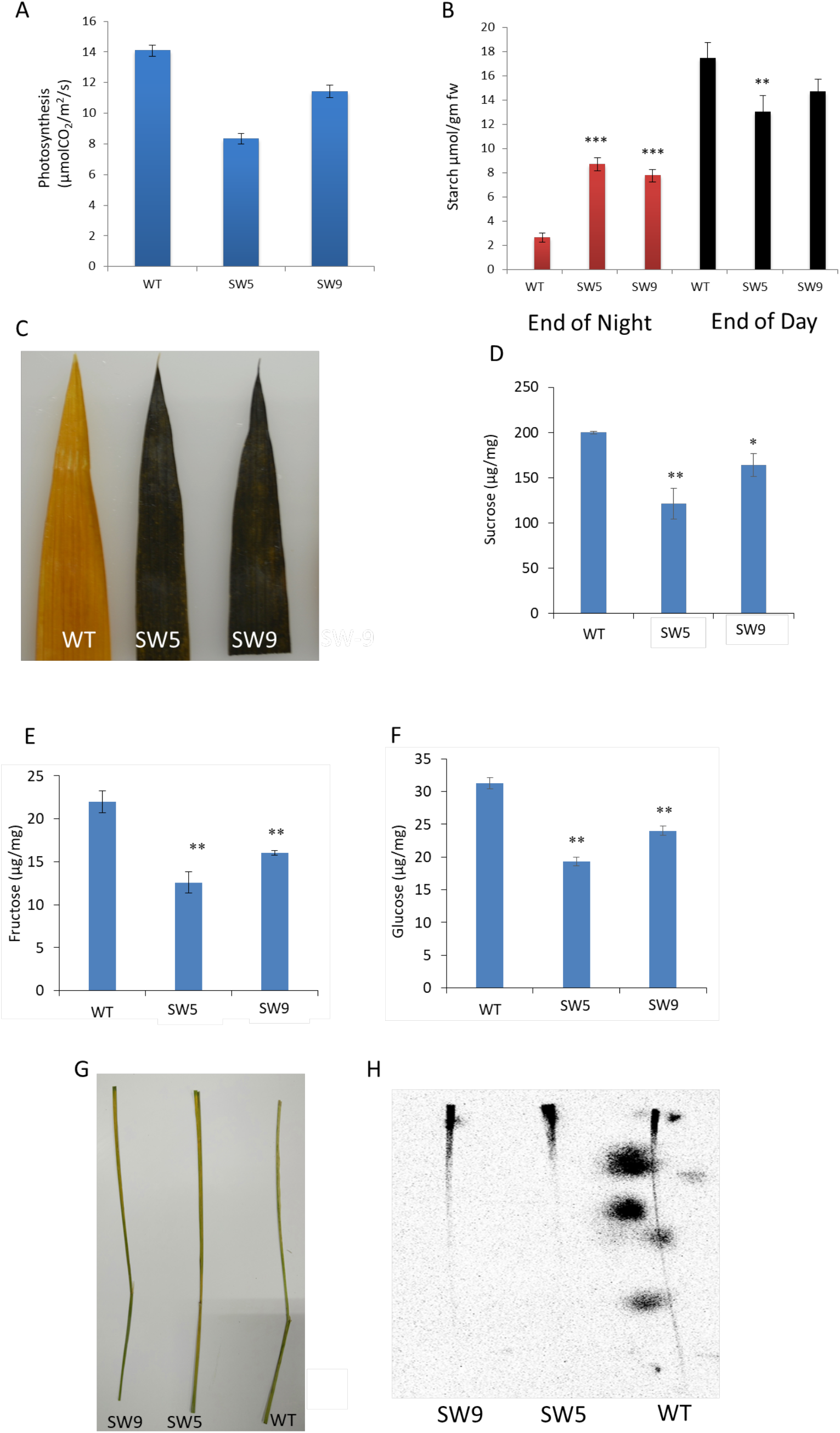
**(A)** Photosynthetic rate in WT and transgenic lines; **(B)** Quantification of starch content in leaf blades of WT, SW5, and SW9 both at the end of night and day; **(C)** Iodine staining of leaf blades at the end of night; **(D-F)** Sucrose, fructose and glucose concentration in leaf blades of WT, SW5 and SW9; **(G)** Radioactivity was applied on the top of the abraded leaf blade; **(H)** Radioactive [^14^C] sucrose translocation assay in leaves of WT and transgenic lines. Error bars show ± SEM. *P ≤ 0.05, **P ≤ 0.01, ***P ≤ 0.001

As OsSWEET11 and 14 are responsible for sucrose transport, soluble sugars content of leaves harvested from WT and transgenics was measured. The sucrose, glucose and fructose content declined by 40%, 43%, and 38% respectively in SW5 whereas the reduction in SW9 was 18%, 27% and 23% respectively, compared to the WT (Figure 2D, E, and F). To directly monitor the movement of sucrose in leaves, radiolabeled [^14^C] sucrose was applied to the top of the leaf blade of an intact plant (Figure 2G). After 1.5 h the radioactive signal penetrated much deeper along the longitudinal axis of the WT leaf. Conversely, in SW5 most of the radioactivity was restricted to the top at the point of its application (Figure 2H). This confirms that the rate of [^14^C] sucrose transport was lower in SW5 than the WT. The SW9 had an intermediate position and its sucrose transport efficiency seemed to be higher than SW5.

Low sucrose translocation also affected growth of roots in transgenic seedlings. Primary roots of SW9 and SW5 seedlings were significantly shorter than the WT when grown in sugar deficient media. Even the shoots of the transgenic seedlings were shorter than the WTs (Figure 3A and D). The results suggested that sucrose translocation from the storage reserves *i*.*e*. seeds to sink tissues was repressed in transgenic seedlings.

**Figure 3.**
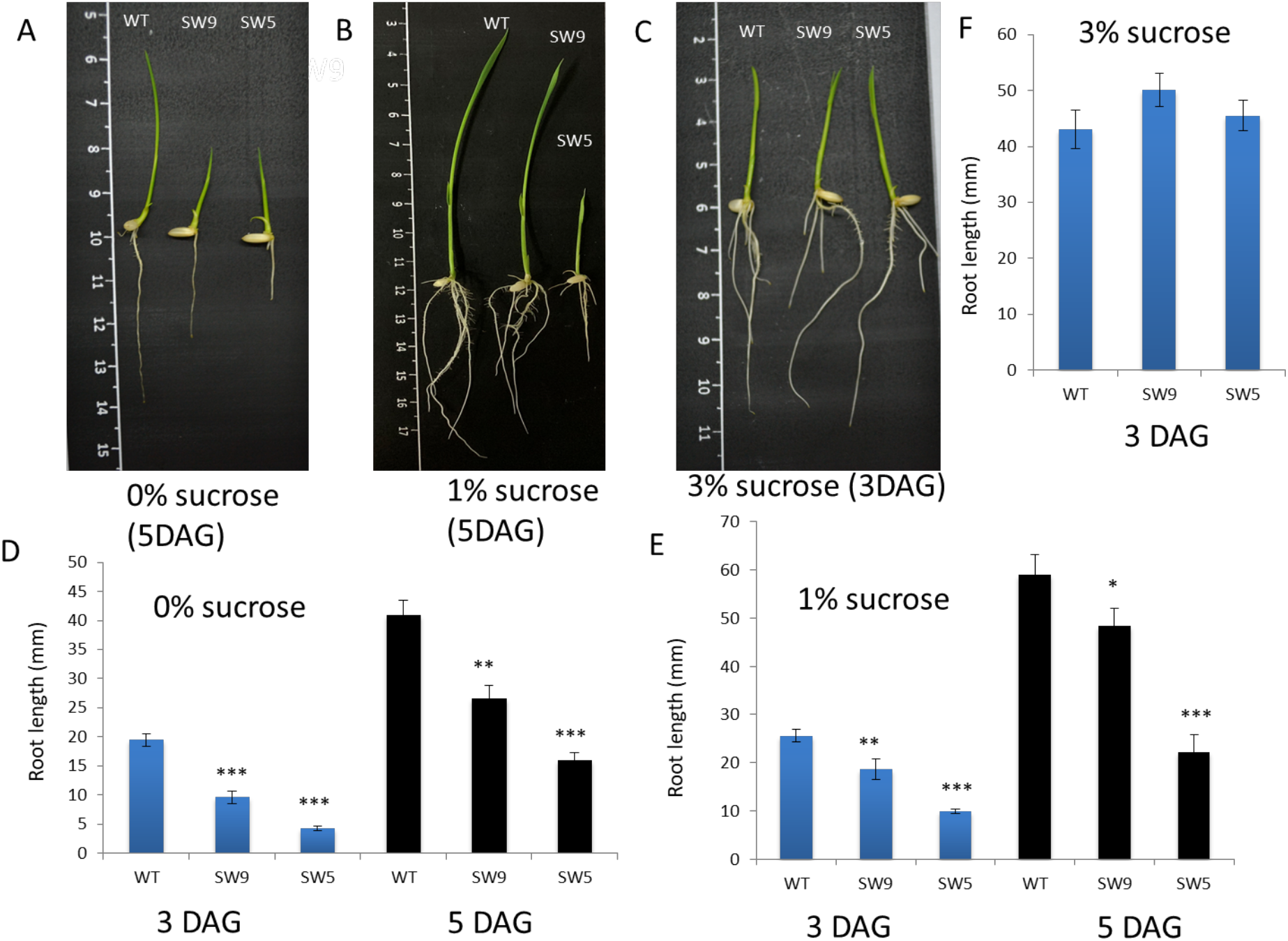
Root growth analysis of WT, SW5 and SW9. **(A-C)** Pictures showing the root and shoot growth of WT, SW9 and SW5 seeds grown on different sugar concentration 5 days after germination (DAG) (0% and 1% sucrose) and 3 DAG (3% sucrose); **(D-F)** Measurement of the primary root length of WT, SW9 and SW5 after 3 and 5 DAG in 0% and 1%, and 5 DAG in 3% sucrose concentration in the growth medium. Medium supplemented with 3% sucrose completely rescued the phenotype of SW9 and SW5 plants **(C and F)**. Blue bars show 3 DAG, black bars show 5 DAG. All the measurements are mean of 3 replicates. Error bars show ± SEM. *P ≤ 0.05, **P ≤ 0.01, ***P ≤ 0.001

### Exogenous sucrose restores normal root and shoot growth in transformed lines

Small rice seedlings up to 7^th^ day of their growth are mostly dependent on carbohydrate reserve of the seed endosperm. Suppressed root and shoot development of the seedlings of transgenic lines was due to insufficient supply of sugars from the endosperm. Seeds of rice transgenic lines grown on media supplemented with 1% sucrose could not rescue the short root and shoot phenotype completely (Figure 3B and E). Growth media with 1% sucrose stimulated root and shoot growth to a larger extent in SW9 as compared to SW5 (Figure 3B and E), suggesting that sucrose transport is inhibited more in SW5 than SW9. However, supplementation of 3% sucrose in growth media completely rescued the short root and shoot phenotype of both SW5 and SW9 seedlings. Transgenic seedlings supplemented with external 3% sucrose grew similar to the WT (Figure 3C and F).

### Expression of genes involved in sucrose synthesis is reduced in transgenic lines

Since the rate of sucrose transport was lower than the WT in SW5 and SW9, we checked the expression of other *OsSUTs* that might contribute to phloem loading and long distance sucrose translocation. The transcript levels of *OsSUT2, OsSUT3, OsSUT4*, and *OsSUT5* in the transgenic lines remained nearly unaffected (Figure 4A). This prompted us to check the expression of genes involved in carbohydrate metabolism in leaves.

**Figure 4.**
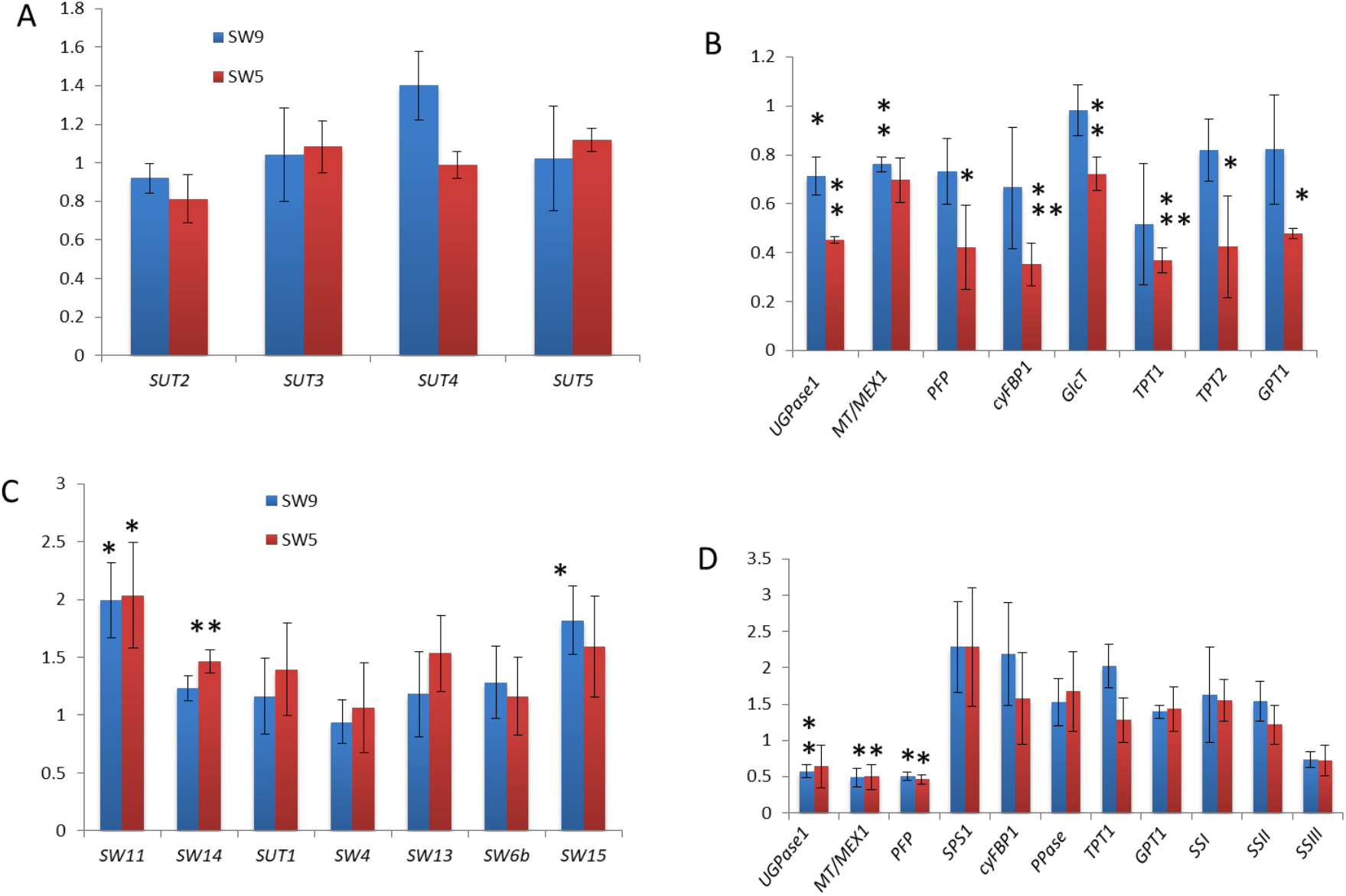
Expression analysis of different genes involved in carbohydrate transport and metabolism by RT-qPCR. **(A and B)** Relative expression levels of different *SUTs* and genes involved in carbohydrate metabolism in leaves; **(C and D)** Relative expression levels of different genes involved in sugar transport, carbohydrate metabolism and grain filling in 6-7 DAP caryopsis. Data shown as mean of 4 biological replicates. Rice *ACTIN1* is used for normalization. Error bars show ± SEM. *P ≤ 0.05, **P ≤ 0.01, ***P ≤ 0.001

Triose phosphates produced by the carbon reduction cycle in chloroplasts are transported to the cytosol by triose phosphate translocators (TPTs) and converted to sucrose by the sequential action of many enzymes such as pyrophosphate fructose-6-phosphate-1-phosphotransferase (PFP), cytosolic fructose-1, 6-bisphosphatase (cyFBP), UDP glucose pyrophosphorylase (UGPase/UDPG), sucrose phosphate synthases (SPSs), etc. The RT-qPCR analysis showed that both SW5 and SW9 transgenic lines had lower expression of *OsTPT1, OsTPT2, OsPFP, cyOsFBP1*, and *OsUGPase1* than the WT (Figure 4B). Transcription of these genes in SW5 was reduced by more than half as compared to the WT (Figure 4B). Lower expression of *OsTPT1* and *OsTPT2* restricted the movement of triose phosphates from chloroplasts, thereby diminishing the substrate concentration for sucrose synthesis in the cytosol of transgenic lines. Downregulation of the genes involved in sucrose synthesis pathway *i*.*e. OsPFP, cyOsFBP1*, and *OsUGPase1* further reduced the rate of sucrose synthesis in transgenics.

Further the transcript levels of maltose transporter (MT or MEX1) and glucose transporter (GlcT) that are responsible for the transport of maltose and glucose, through the transitory starch-mediated salvage pathway were reduced in transgenic lines in comparison to WT (Figure 4B). Lower photosynthetic rates and decreased expression of genes involved in sucrose synthesis and membrane transporters *TPT1, TPT2, MT*, and *GlcT* in transgenic lines resulted in severe shortage of soluble sugars such as sucrose, glucose, and fructose, in leaves of transgenic plants (Figure 2D-F).

### Transgenic lines have aberrant grain filling

The SW5 and SW9 transgenic lines had aberrant grain filling leading to chalkiness, shorter, compressed and wrinkled seeds (Figure 5A-D). Approximately 30% of the seeds in both the transgenic lines were abnormal; line SW5 being severely affected. Wrinkled appearance of the grains of transgenic lines SW5 and SW9 was apparent 6-7 days after pollination (DAP) (Figure 6A). SW5 grains were more compressed than the SW9. Grains from both SW9 and SW5 lines had shorter diameter than the WT (Figure 6A). Consequently, the average weight of 100 seeds declined in both the transgenic lines (Figure 1H).

**Figure 5.**
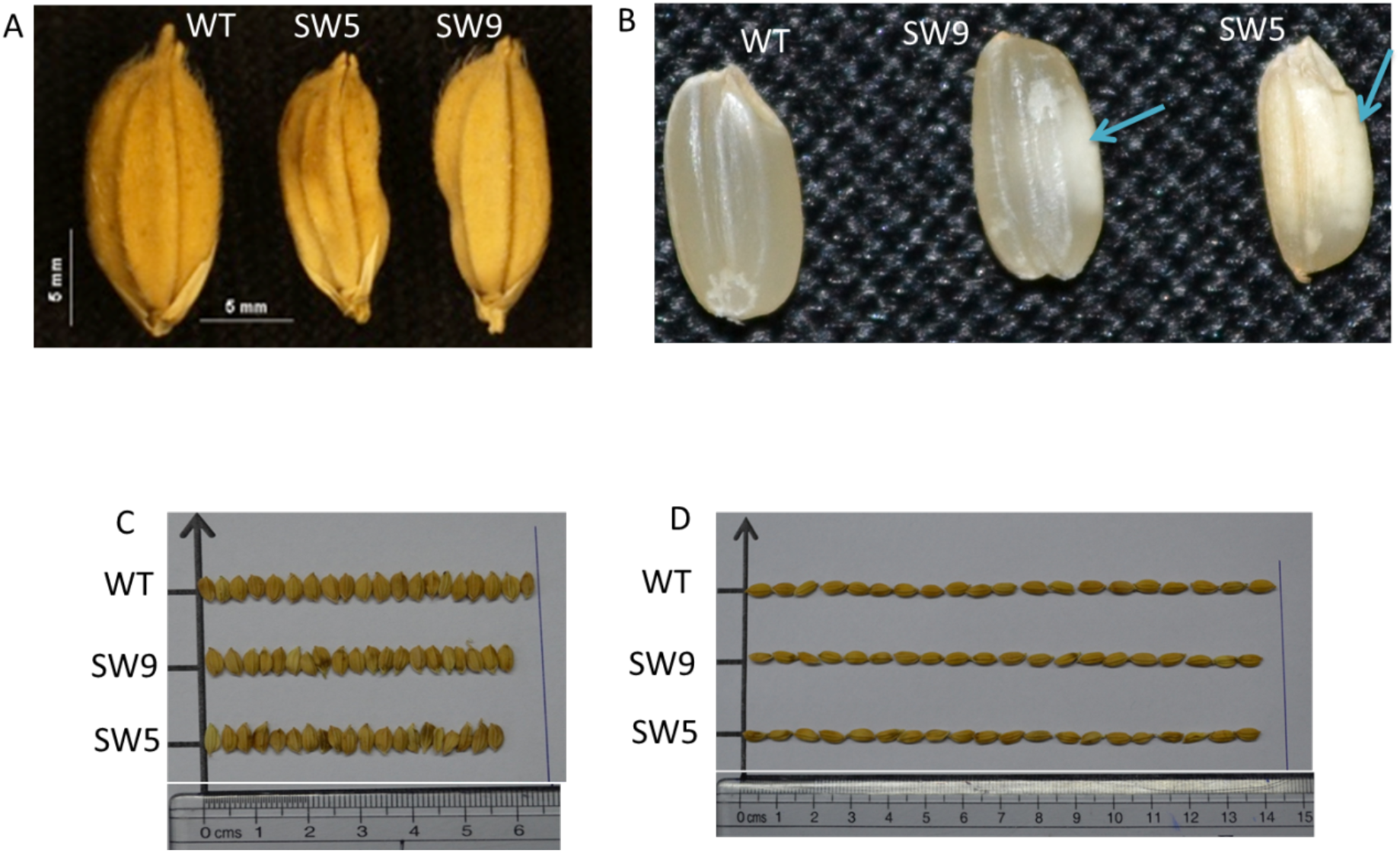
Morphology of seeds. **(A)** Wrinkled and shorter grains of lines SW5 and SW9 as compared to the WT; **(B)** Improper grain filling in transgenic lines leading to chalkiness (blue arrows indicating white belly); **(C and D)** Width and length of 20 seeds selected randomly from WT, SW9 and SW5 plants.

**Figure 6.**
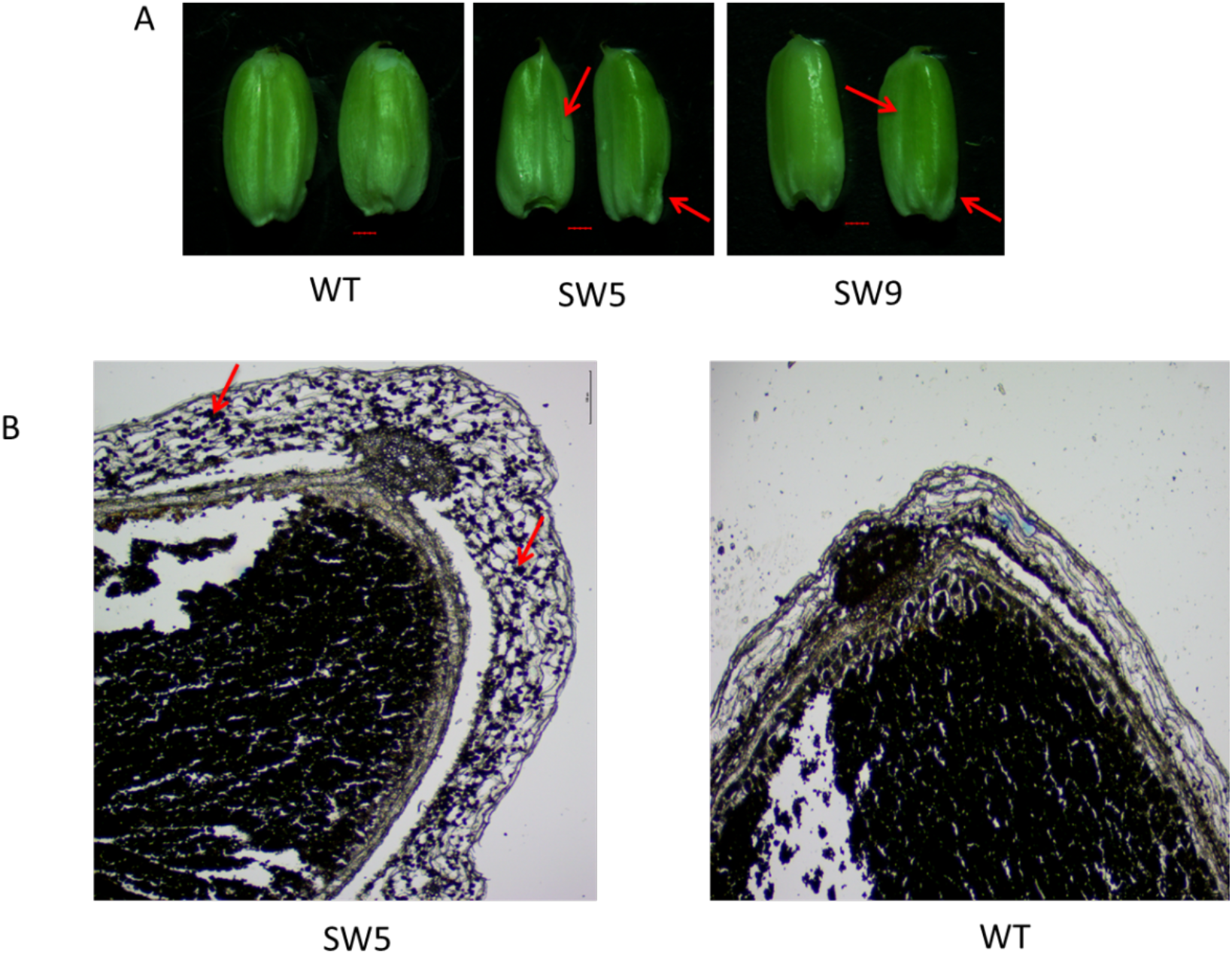
**(A)** Morphological defects in seeds of WT and transgenic lines appeared 6-7 DAP; **(B)** Histological analysis of 9 DAP old grains. Iodine staining shows starch accumulation in the pericarp of SW5 indicating abnormal sugar metabolism and grain filling in transgenic lines. Bar in (A) is same for WT, SW5 and SW9. Bar is 100µm (B).

Seven to nine days after pollination (DAP) transient starch in the pericarp is usually transported to the endosperm (Wu et al., 2016). Histological analysis of 8-9 DAP old caryopsis showed high accumulation of starch in the pericarp of SW5 as compared to the WT (Figure 6B). Presence of starch granules in the pericarp of SW5 even 9 DAP indicates that sucrose transport from pericarp to the endosperm is affected resulting in accumulation of starch in the pericarp.

### Expression of genes involved in sucrose-starch metabolism is altered in seeds of transgenic lines

Defect in the grain morphology of our transgenic lines was quite apparent after 6-7 DAP (Figure 6A). RT-qPCR analysis of 6-7 DAP caryopsis showed that the expression of *OsSWEET11, OsSWEET14*, and *OsSUT1* was increased in transgenic lines SW5 and SW9 in comparison to the WT; with SW5 having higher transcript levels than SW9 (Figure 4C). Transcriptional activity of other sugar transporters expressed during grain filling *i*.*e. OsSWEET13, OsSWEET15, OsSWEET4* and *OsSWEET6a*, was either unchanged or slightly increased in both SW5 and SW9 lines compared to the WT (Figure 4C). As in case of leaves, the expression of sugar metabolizing genes, *OsUGPase1, OsMT* and *OsPFP1*, was reduced by half in transgenic lines SW5 and SW9 as compared to WT (Figure 4D). It is apparent that the incomplete grain filling in lines SW5 and SW9 is due to impaired carbohydrate metabolism and transport in their caryopses as well as leaves.

### Transgenic lines were less susceptible to *Xoo*

As the rice bacterial blight pathogen *Xoo* is known to co-activate *OsSWEET14/Os-11N3* along with *OsSWEET11/Os-8N3/Xa13*, overexpression of both these genes is expected to enhance the rice susceptibility against *Xoo*. However, in a virulence assay, 10 days post leaf clip inoculation with *Xoo* strain PXO99^A^, leaves of SW9 and SW5 had less susceptibility to infection as compared to WT leaves (Figure. 7A). WT Nipponbare plants had an average of ∼ 6.5 cm lesions, whereas both SW9 and SW5 transgenic plants showed significantly lower lesion lengths of approximately 3.5 cm and 0.8 cm respectively (Figure. 7B). This indicates that the resistance to pathogen attack was higher in SW5 and SW9 transgenic lines than the WT. However, complete leaf yellowing was observed both in WT and transgenic lines after longer infection times (*i*.*e*. one month post inoculation).

**Figure 7.**
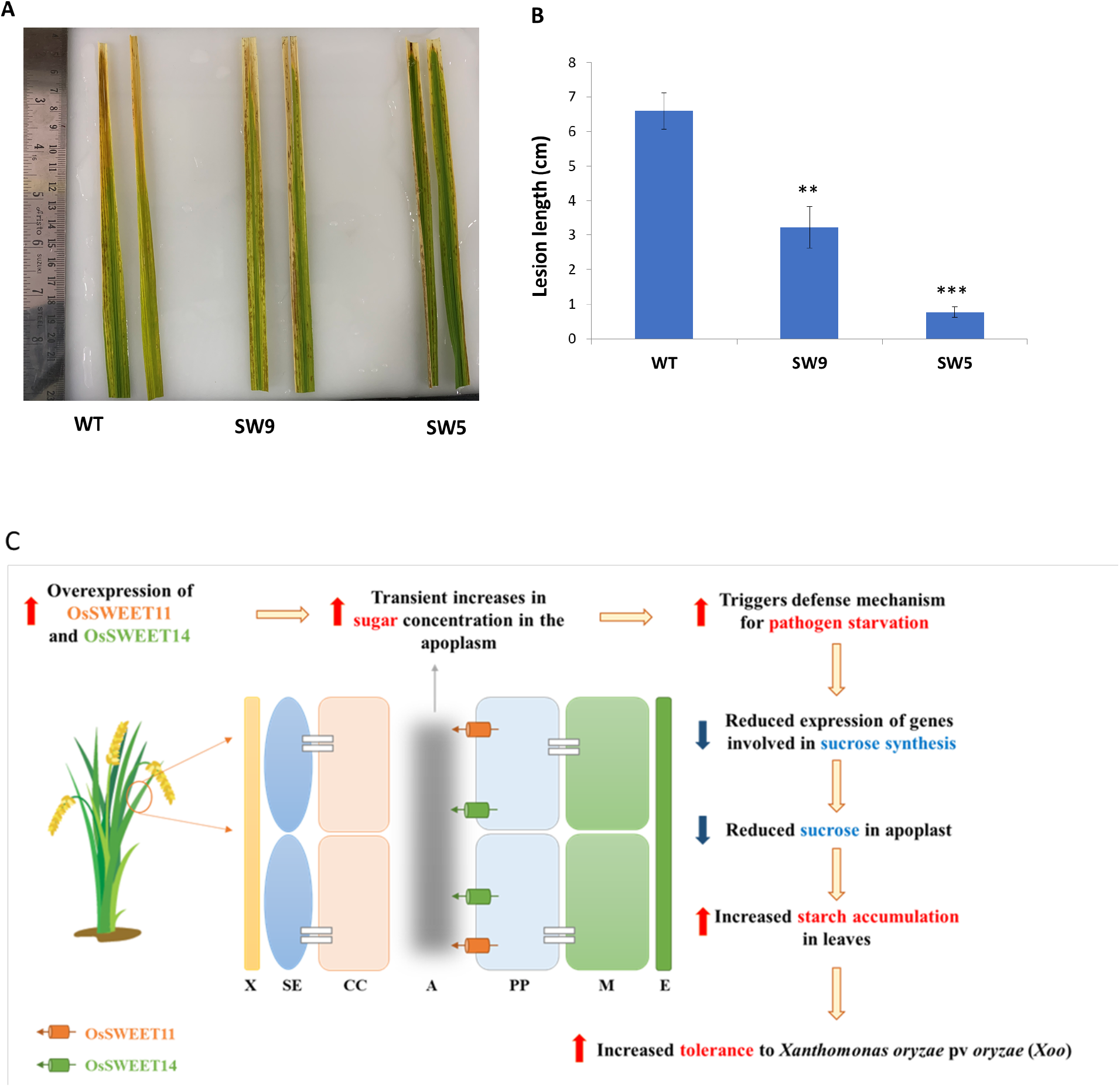
Virulence assay with *Xoo* strain PXO99^A^. **(A)** Images of leaves of transgenic lines WT, SW9 and SW5, 10 days post leaf clip inoculation with *Xoo* strain PXO99^A^ **(B)** Lesion lengths in transgenic and WT plants 10 days post infection. **(C)** Pictorial representation of the likely changes occurring in transgenic plants due to the enhanced expression of *OsSWEET11* and *OsSWEET14*. Activation in expression of OsSWEET11 and OsSWEET14 might lead to transient increase in apoplasmic sugar concentration, which is sensed as a pathogen (*Xoo*) attack by the plant. This triggers the plant defense mechanism for pathogen starvation. Drastic reduction in the expression of genes involved in transport of triose phosphates from chloroplast and sucrose synthesis (refer to text) takes place leading to decreased concentration of sucrose in the apoplasm and hyper accumulation of starch in chloroplasts. Low levels of sucrose in the apoplasm act as a deterrent for *Xoo* growth and thus contribute to the low susceptibility of transgenic lines. Error bars show ± SEM. *P ≤ 0.05, **P ≤ 0.01, ***P ≤ 0.001. **X**-xylem, SE-sieve elements, **CC**-companions cells, **A**-apoplasmic space, **PP**-phloem parenchyma, **M**-mesophyll cells, **E**-epidermis.

## Discussion

### Tissue specific co-expression of *OsSUT1, OsSWEET11*, and *OsSWEET14* hampers plant growth and yield

A recent study has shown that overexpression of *OsSUT1* in rice leads to increased seed size and vegetative growth (Feng et al., 2018). Similarly, ectopic expression of *AtSUC2*, the ortholog of rice *SUT1* in *Arabidopsis*, hooked to companion cell specific *PHLOEM PROTEIN 2* promoter resulted in improved phloem loading in rice (Wang et al., 2015). Enhanced sucrose loading and translocation led to increased biomass and yield than the WTs under field conditions (Wang et al., 2015). However, constitutive overexpression of *OsSWEET11* or *OsSWEET14* in rice resulted in stunted plants and reduced yield as compared to WT (Gao et al., 2018; Kim et al., 2020). Similarly, transgenic *Arabidopsis* plants showed growth retardation when orthologs of *OsSWEET11* and *OsSWEET14* in *Arabidopsis i*.*e. AtSWEET11* and *AtSWEET12* respectively, were constitutively expressed (Eom et al., 2015). SWEETs mediated sucrose transport in plants occur in specialized cells like vascular parenchyma and mesophyll cells of leaves and tissues of developing grains like nucellar projection, nucellar epidermis etc. (Eom et al., 2015; Yang et al., 2018). Therefore, constitutive and ubiquitous expression of SWEETs could result in loss of directionality leading to random and futile movement of sucrose molecules across cells causing unintended effects (Eom et al., 2015). As OsSUT1, OsSWEET11, and OsSWEET14 work in co-ordination, and are co-regulated by OsDOF11, it was plausible that global increase in the expression of OsSWEET11 or OsSWEET14 alone desynchronized their functioning that led to imbalances in sugar partitioning and transport in rice.

Therefore, we expressed *OsSUT1, OsSWEET11*, and *OsSWEET14* under their native promoters to maintain their cell specific expression and synchronized movement of sucrose in all tissues. Our transgenic lines had significantly higher expression of *OsSUT1, OsSWEET11*, and *OsSWEET14* than the WT. Stable and transient transgenic tobacco transcriptional lines of the *pOsSWEET11-GUS* and *pOsSWEET14-GUS* constructs, respectively, were generated to test the tissue specific expression of these genes. GUS expression only in leaf vein network was observed in both the cases (Figure S3). This indicates that *OsSWEET11* and *14* in our transgenics were expressed appropriately in tissues where they are indigenously present. Similar pattern of GUS expression is observed when *GUS* hooked to promoters of *AtSWEET11* and *AtSWEET12*, is expressed in transgenic *A. thaliana* plants (Chen et al., 2012). We found that even the tissue specific concurrent overexpression of *OsSWEET11* with *OsSUT1* and *OsSWEET14*, adversely affected plant growth and yield as observed in transgenic rice plants overexpressing *OsSWEET11* and *OsSWEET14* (Gao et al., 2018; Kim et al., 2020).

Transcription of *OsSUT1, OsSWEET11*, and *OsSWEET14* is co-regulated by transcription factor OsDOF11. It has been shown that constitutive overexpression of the transcription factor *OsDOF11* in rice did not affect plant growth (Wu et al., 2018). Transcript levels of *OsSUT1, OsSWEET11*, and *OsSWEET14* were not changed significantly in *OsDOF11* overexpressor plants as compared to the WT (Wu et al., 2018). However, in another very recent study plant height of two out of the 4 OsDOF11 overexpressing lines appeared smaller than the WT and all the overexpressor lines had reduced grain yield as compared to the WT (Kim et al., 2020). In this case expression of *OsSWEET14* was increased substantially in OsDOF11 overexpressor in comparison to WT (Kim et al., 2020). Tissue specific transcriptional activation of OsDOF11 by fusion of VP16 resulted in increased 1000 grain weight of transformed rice lines as compared to WT, possibly through stimulating *OsSWEET14* expression (Kim et al., 2020). However, yield increments in transformed lines were not high.

### Impaired carbohydrate metabolism and transport in leaves and seeds led to phenotypic defects

Despite increased native expression of *OsSUT1, OsSWEET11* and *OsSWEET14*, our transgenic lines exhibited phenotypes of impaired sucrose synthesis and transport such as deposition of starch in leaves, reduced plant height, wrinkled and floury grains etc. Decrease in the rate of sugar synthesis and its reduced export out of chloroplast was due to drastically reduced expression of genes required for transport of triose phosphates and sucrose biosynthesis *i*.*e. OsTPT1, Os*TPT2, *OsPFP, cyOsFBP1*, and *OsUGPase1*, in the leaves of transgenic lines (Figure 4B). Triose phosphates exported from the chloroplast to the cytosol via TPT1 and TPT2 are substrate for sucrose synthesis in cytosol. *OsTPT1* is the predominant gene expressed in the chloroplast envelope of photosynthetic leaves. Rice mutants, *ostpt1-1* and *oscfbp1*, that had reduced availability of triose phosphates and fructose-6-phosphate, respectively, in the cytosol as substrates for sucrose synthesis had stunted growth phenotype (Lee et al., 2008; Lee et al., 2014). Rice mutants of *ugpase1*, also referred to as *flo8*, too were shorter than the WTs (Long et al., 2017). In our transgenic lines we found a significant reduction in the transcript levels of genes involved in sucrose transport and biosynthesis, including *OsTPT1, OsUGPase1*, and *cyOsFBP1*. The *OsTPT1* and *cyOsFBP1* showed highest down regulation in both SW5 and SW9 lines (Figure 4B). Consequently, the content of sucrose and other soluble sugars, fructose and glucose, was considerably reduced in transgenic lines as compared to WT (Figure 2D-F). This was further compounded by the lower photosynthetic rates in transgenic lines causing a reduction in carbon fixation. These downregulated the vegetative growth and developmental processes of rice that resulted in decreased plant height, biomass and yield.

Conversely, transgenic/mutant plants with decreased/nil expression of *TPT1* and *cyFBP* in dicot species such as *Arabidopsis*, tobacco and potato, did not affect plant growth under ambient light and CO_2_ conditions (Lee et al., 2008). Reduced supply of sucrose in these mutants is compensated by a concomitant increase in the starch content of leaves. This starch mediated salvage pathway often provides substrates for sucrose synthesis (Lee et al., 2008). Starch is primarily degraded in chloroplasts to maltose by the action of β amylase and isoamylase3 (Zeeman et al., 2004). The other product of starch degradation is glucose. Maltose and glucose are transported via MT and plastidic glucose translocator (OspGLcT) respectively, to the cytosol for sucrose synthesis. In rice, starch breakdown pathway is not sufficient to replenish the decreased sucrose levels in the cytosol to sustain normal plant growth. Majority of the assimilated carbon flux (more than 85 %) is directed to soluble sugars in rice compared to 54-60% in *Arabidopsis* (Lee et al., 2014). In transgenic lines SW5 and SW9, the expression levels of *OsGLcT* and/or *OsMT* was lower than the WT (Figure 4B). This further diminished any possibility of circumventing the sucrose shortage in cytosol through starch degradation pathway. Decreased expression of *OsMT* and *OsGlcT* and the restricted movement of triose phosphates led to hyper accumulation of starch at the end of dark period in leaves of transgenic lines. Further, due to reduced vegetative growth of transgenics that needed lower energy, the rate of respiration would have declined leading to the accumulation of starch at the end of the night. Leaves of *Arabidopsis* mutants of *mex1* also accumulate excess starch (Niittylä et al., 2004). Accumulation of carbohydrates in leaves either by removing active sink tissues or impairing sucrose transport from source leaves generally leads to photosynthesis inhibition (Ainsworth and Bush, 2011). This could be the reason for lower photosynthetic rates in SW5 and SW9 than the WT.

Overexpression of *OsSUT1* and *AtSUC2* (Arabidopsis ortholog of *OsSUT1*) in rice plants has been shown to increase grain width and yield respectively (Wang et al., 2015; Feng et al., 2018). However, transgenic rice plants constitutively expressing *OsSWEET11* under *UBIQUITIN 1* promoter had reduced yield than the WT (Gao et al., 2018). Similarly, seeds of our transgenics lines also had reduced yields with multiple morphogenic defects such as incomplete filling, deformation, and chalkiness. As in case of leaves, RT-qPCR analysis revealed severe reduction in expression of *MT, UDPG1/ UGPase1*, and *PFP* genes involved in sucrose synthesis in the pericarp of developing rice caryopsis causing deposition of starch granules in the pericarp (Figure 4D and 6B).

*UGPase1/FlO8* and *PFP* are vital for the normal seed development in rice (Duan et al., 2016; Long et al., 2017; Chen et al., 2020). Inactivation by RNAi or mutation in *ugpase1/flo8* gene in rice plants causes chalkiness in the grains with floury endosperm that is also observed in our transgenic lines (Woo et al., 2008; Long et al., 2017). Its suppression results in male sterility and abnormal callose deposition (Chen et al., 2007). No symplasmic movement of sugar molecules takes place in rice pollens due to the absence of plasmodesmatal connection to the neighboring sporophytic tissue. UGPase1 provides an apoplasmic sugar unloading pathway for pollen development by providing sucrose (Chen et al., 2007). Similarly, no symplasmic pathway for sugar transport to maternal filial tissues from the surrounding sporophytic cells is reported in rice and thus the presence of sugar transporters is inevitable (Yang et al., 2018). UGPase1 forms sucrose from the transitory starch in the pericarp, which is translocated to the filial cells by these transporters via the apoplasmic space between endosperm and embryo (Ludewig and Flügge, 2013). Further, in endosperm of cereals such as barley the activity of UGPase is required to produce Glu-1-P from UDP-Glucose (Kleczkowski, 1994). Glu-1-P is converted to ADP-Glucose, the substrate for starch synthesis in endosperm, by the action of AGPase (Kleczkowski, 1994). Therefore, reduced expression of *UGPase1* in our transgenic lines might have led to sucrose shortage in the pericarp for grain filling by the sugar transporters and simultaneously compromised the starch biosynthesis in seed endosperm by limiting Glu-1-P levels.

PFP catalyzes the reversible conversion of fructose-1, 6-bisphosphate to fructose-6-phosphate and can therefore shift the flux towards either gluconeogenesis, resulting in sucrose synthesis, or glycolysis, leading to pyruvate production. Its expression during both early and late stages of seed filling in rice is important for seed development (Duan et al., 2016). Mutation in rice *pfp* has been shown to affect carbon metabolism in seeds that results in aberrant grain filling. Rice *pfp* mutants had chalkiness in seeds, reduced kernel thickness and grain weight (Duan et al., 2016; Chen et al., 2020). Severe downregulation of *PFP* in our transgenic lines (Figure 4D), would have impaired the carbon metabolism in seeds by diminishing the provisions of sucrose formation from transiently stored starch leading to its accumulation in the pericarp.

OsMEX1/MT is the rice maltose transporter present in plastidial inner membrane. Expression profiling of *OsMEX1* revealed that it is expressed in sink tissues such as pollen grains, flowers, seeds and panicles (Ryoo et al., 2013). MEX1 expression in sink tissues of different species such as rice, *Arabidopsis*, and apple are indicative of its role in starch mobilization in sink tissues (Reidel et al., 2008; Ryoo et al., 2013). More than 50% reduction in the transcript levels of *OsMEX1* in seeds of our transgenic lines as compared to WT (Figure 4D), and the associated phenotype of abnormal grains (Figure 5A and B) further strengthens the possibility of *OsMEX1* being a key player in assimilate partitioning in sink tissues. Generation of knockout lines of *OsMEX1* could be useful to confirm its role in starch metabolism and sugar transport in sink tissues such as seeds. Therefore, the aberrant grain filling in our transgenic lines is a result of impaired sucrose and starch mobilization due to strong down regulation of *OsUGPase1* and *OsPFP*, along with *OsMEX1*.

### Overexpression of *OsSWEET11* and *14* modulates plant immunity against *Xoo*

*Xa13/Os8N3/OsSWEET11* and *Os11N3/OsSWEET14* were initially discovered as host susceptible genes that are induced directly upon infection by *Xoo* for its survival (Chu et al., 2006; Yang et al., 2006). Our virulence assay shows that the transformed lines SW5 and SW9 overexpressing *OsSWEET11, OsSWEET14* and *OsSUT1* are less susceptible to *Xoo* as compared to WT (Figure. 7A). Overexpression of *OsSWEET11* increased the susceptibility of transgenic rice plants to both *Xoo* and rice sheath blight pathogen *Rhizoctonia solani* (Gao et al., 2018). Conversely, transgenic rice plants overexpressing *OsSWEET14* are found to be less susceptible to *R. solani* due to reduced sugar concentration in the apoplasm (Kim et al., 2020). Similarly, overexpression of OsDOF11 that regulates the expression *OsSUT1, OsSWEET11*, and *OsSWEET14*, in transgenic rice plants, improved resistance to *R. solani* via activation of *OsSWEET14*, which in all probability reduced the sugar content in the apoplasm (Kim et al., 2020). However, sugar metabolism and concentration of soluble sugars in leaves were not analyzed in any of the studies.

Our results demonstrate that transgenic rice lines overexpressing *OsSUT1, OsSWEET11*, and *OsSWEET14* were less susceptible to *Xoo* due to reduced concentration of soluble sugars in leaves of transgenics as compared to WT (Figure 2D-F). We believe that as hypothesized in the sugar signalling model of pathogen resistance (Bezrutczyk et al., 2018), upregulation of rice SWEETs, particularly, SWEET14, result in transient increase in the apoplasmic sugar content, which is perceived as a signal of *Xoo* attack. Upregulation of SWEETs could elicit host defense responses via translocation of sugars to sites of infection (Bezrutczyk et al., 2018). Activation of SWEETs leading to the downregulation of genes involved in sucrose synthesis and restriction of its export from leaf photosynthetic cells to the apoplasm by storing it as starch in chloroplasts might be one of the strategies to limit the quantity of sugars in the extracellular space to inhibit pathogen growth (Figure. 7B), as contemplated in the ‘pathogen starvation’ model of disease resistance in plants (Bezrutczyk et al., 2018). Sugar transporters such as proton hexose symporters that take up hexoses from apoplasmic space to the cytoplasm, are also induced in *Arabidopsis* during pathogen challenge to lower the sugar concentrations in apoplasm and consequently inhibiting pathogen growth (Lemonnier et al., 2014; Yamada et al., 2016). Thus, it seems that upregulation of SWEETs triggers both the sugar signalling and pathogen starvation defense models, and rice plants strike a balance between sugar transport and pathogen resistance by maintaining a steady state level of SWEETs in the cell. Enhancing sucrose transport by inducing *SWEET11* and *14* might put the plant at a disadvantage by making it susceptible to pathogen attack. This could be the reason for stunted growth of transgenic rice plants overexpressing *OsSWEET11* and *OsSWEET14* (Gao et al., 2018; Kim et al., 2020), and *Arabidopsis* plants transformed with *AtSWEET11* and *12* (Eom et al., 2015).

*Xoo* strains can overcome plant defenses by modulating the host expression of genes and suppressing the defense machinery by secreting effectors (Jha and Sonti, 2009). Hundreds of genes from various pathways including glycolysis and sucrose synthesis are differentially upregulated in leaves of susceptible rice variety JG30 infected with PXO99^A^ as compared to JG30 infected with mutant PXO99^A^ without TAL effector *i*.*e*. PH (Tariq et al., 2019). The expression of *UGPase, PFP* and *FBP*, the genes associated with sucrose metabolism, was significantly increased in JG30 infected with, PXO99^A^ than PH, hinting towards their role in pathogen virulence and survival (Tariq et al., 2019). In our transgenic lines *OsSWEET11* and *OsSWEET14* were upregulated, but genes of sucrose synthesis pathway including *OsUGPase1, OsPFP1* and *OsFBP1*, and its transport were downregulated leading to reduced sugar concentrations in the apoplasm. The decreased expression of these genes in our transgenic lines could be a plant defense strategy to reduce sucrose concentration in apoplasm, which otherwise would have been suppressed in the presence of TAL effectors secreted by PXO99^A^. On the basis of our findings we have proposed a probable model of the sequence of metabolic and physiological events that occur due to the enhanced expression of *OsSWEET11* and *OsSWEET14* leading to the phenotypic defects in rice plants and pathogen resistance (Figure 7B).

## Methods

### Vector construction and generation of transgenic rice

Full length sequence of *OsSWEET11* and *OsSWEET14* containing the promoter region and the CDS (introns and exons) was PCR amplified using gDNA of rice (*Oryza sativa* L. ssp *japonica* cv Nipponbare) with sequence specific primers. The promoter and CDS of OsSUT1 were amplified separately from gDNA and cDNA respectively with sequence specific primers. *CaMV 35S* terminator sequence was amplified from pGreenII 0029 62-SK. All the amplified products were initially cloned into pCR-4-TOPO vector (Invitrogen, USA). All the sequences were retrieved from NCBI or Rice Genome Annotation database. Individual expression cassettes of *OsSWEET11, OsSWEET14* and *OsSUT1* with their native promoters and *CaMV 35S* terminator were made in gateway compatible entry vectors EV-1 (pL12R34-Amp) and EV-2 (pL12R34) by restriction digestion and ligation. *OsSWEET11* and *OsSWEET14* fragments were cloned directionally in *BamH1* and *Not1* sites, of EV-1 and EV-2 respectively. *OsSUT1* promoter was cloned in *Xho1* and *BamH1* sites, whereas *OsSUT1* cds was cloned in *BamH1* and *Not1* sites of EV-1. *CaMV 35S* terminator sequence was cloned in *Not1* and *Sac1* sites of EV-1 and EV-2. MCS of EV-1 and 2 is shown in figure S1. All three gene cassettes were pyramided into a single plant transformation vector *pMDC99* (containing *HPTII* gene as plant selection marker) using the LR recombinase mediated MultiRound-gateway cloning process (Chen et al., 2006; James et al., 2018). The recombinant *pMDC99-OsSWEET11;OsSWEET14;OsSUT1* plant transformation vector was transformed into *Agrobacterium* EHA105 and was used for transformation of *japonica* rice (*Oryza sativa* cv Nipponbare). Transgenic plants were raised according to the protocol followed in our previous study (James et al., 2018). Sequence of all PCR primers used in the study is mentioned in **Table S1**.

### Plant growth conditions and photosynthesis measurement

T_2_ or T_3_ homozygous lines of transgenic plants were grown in pots in green house with 16hr/8hr day/night conditions. Tiller number, plant height, panicle number, panicle length, flag leaf (associated with the highest panicle) width measurement and seed collection were done from plants grown in green house. For measurement of root length, seeds of WT and transgenic lines were grown in ½ MS media supplemented with desired concentration of sucrose. Root length is measured using ImageJ software. Photosynthetic rate was measured using an infrared gas analyzer (Licor 6400-XT portable photosynthetic system) as described previously (Kandoi et al., 2016), with following chamber conditions, PAR-1000 µmol photons/m^2^/s, [CO_2_]-400ppm, Temperature-30°C, and 55% relative humidity.

### Identification and validation of transgenic rice lines

Putative transgenic lines were screened by gDNA PCR for the presence of recombinant cassette. Gene specific forward and *CaMV 35S* terminator reverse primers for the transformed genes (*OsSWEET11, OsSWEET14*, and *OsSUT1*) and *HPTII* specific primers were used for PCR (Table S1).

The PCR positive transgenic lines were further confirmed by Southern hybridization of *HPTII* gene with DIG-labelled gene probes (James et al., 2018). For post-hybridization and signal detection the procedure mentioned in the kit was followed (Roche Diagnostics, Germany).

### Leaf GUS assay

Stable transgenic tobacco (*Nicotiana tabacum* cv. Xanthi) lines transformed with transcriptional fusion reporter construct *pOsSWEET11:GUS* was generated as described earlier (Sanan-Mishra et al., 2005), using *Agrobacterium tumefaciens* (LBA4404). Transient GUS expression assay with *pOsSWEET14:GUS* was done as previously described (Lee and Yang, 2006) with little modifications. Promoter of 1500bp for both *OsSWEET11* and *OsSWEET14* was cloned in *BamH1* and *EcoR1* sites of pCambia 1391z vector. Leaf GUS staining was done as described previously (Sanan-Mishra et al., 2005).

### Sugar and starch estimation

For starch estimation the upper portion of the second longest leaf from the tip measuring 8-9 cm (weighing between 0.15 to .25gm) was harvested from WT and transgenic plants at the end of the day and night period. Leaf samples were boiled in 80% aqueous vol/vol ethanol for 3 min. The pellet fraction after centrifugation was used to measure the starch content in the leaf samples by recording the reduction of NAD to NADH in an enzymatically coupled reaction in a spectrophotometer (Smith and Zeeman, 2006).

Frozen leaves (100 mg) were homogenized in liquid nitrogen using mortar and pestle and 1,400 µl of 100% ice-cold methanol (HPLC grade, Sigma) was added. The mixture was vortexed for 30 sec followed by addition of 60 µl of ribitol (0.2 mg/ml) as an internal standard (Lisec et al. 2006). Further processing of samples was done as previously described (Tanna et al., 2018). Methoxyamine hydrochloride was used as derivatizing agent. Derivatized samples were analyzed by a gas chromatography coupled with mass analyze (GC-MS QP-2010, Shimadzu, Japan) equipped with an autosampler (AOC-5000, Shimadzu, Japan) using RTX-5 fused silica capillary column. The mass spectra were recorded with mass range from 45 to 700 m/z. The metabolites were identified by their mass fragmentation and compared with the NIST library and quantified using internal standard ribitol. All the conditions were kept same as described earlier (Tanna et al., 2018).

### Quantitative PCR analysis

RNA was isolated from the longest leaf blade of vegetatively mature plants and also from 6-7 DAP caryopsis of transgenic and WT plants (Yin et al., 2016). For RT-qPCR random hexamers were used for cDNA preparation. CaMV 35S terminator primer was used for first strand cDNA synthesis for RT-PCR (Verso cDNA synthesis kit). RT-PCR using gene specific forward and *CaMV 35S* terminator reverse was performed for expression analysis of transgenes in the WT and transgenic lines. Both RT-qPCR and RT-PCR were performed using standard protocols.

### Radioactive sucrose transport assay

The second leaf from top of each line was selected for this assay. The D-Sucrose-U-14C (0.1 mCi from BRIT, DAE, Govt. of India) was diluted to 0.4µCi in a solution of 4.5 mM unlabeled sucrose. Two microliter of this solution was applied thrice at an interval of 30 minutes to the abraded region (10 cm above the ear on the adaxial side) of each leaf (Wu et al., 2018) After 90 min. leaves were exposed to a phosphor screen and scanned by an image analyzer (Typhoon 9210, GE Healthcare).

### Histological analysis

Caryopsis (8-9 DAP) of SW5 and WT were fixed in 4% Paraformaldehyde in 100mM sodium phosphate buffer (pH 7.4), vacuum infiltrated and incubated overnight at 4 °C. Dehydration was done with increasing ethanol concentration series followed by embedding in paraplast plus (Hirose et al., 2002; Yang et al., 2018). Sectioning was performed with rotary microtome Leica RM 2265. Sections were deparaffinized using xylene and stained with Lugol’s solution for starch staining.

### Virulence assay

Leaf clip inoculation assay was performed as per the protocol given previously (Ke et al., 2017).

### Statistical analysis

A two tail paired student’s t-test was done to check statistically significant differences between values of WT, SW5, and SW9 samples in each experiment. SW8 was included in some experiments as mentioned.

## Acknowledgements

JS acknowledges the grant provided by University grants commission Govt. of India under Dr. D S Kothari Post Doctoral Fellowship scheme. The authors acknowledge the administrative and logistical support from Jawaharlal Nehru University and National Institute of Plant Genome Research, New Delhi, India.

## Authors’ contributions

J.S. and B.C.T. conceived the project idea and designed experiments. J.S., D.J., V.M.M.A. and M.K.P. performed the experiments. J.S., B.C.T., M.K.R. and J.K.T. analyzed the data. J.S. and B.C.T. wrote the manuscript.

## References

Ainsworth, E. A., and Bush, D. R. (2011). Carbohydrate export from the leaf: A highly regulated process and target to enhance photosynthesis and productivity. Plant Physiol. Advance Access published 2011, doi:10.1104/pp.110.167684.

Antony, G., Zhou, J., Huang, S., Li, T., Liu, B., White, F., and Yang, B. (2010). Rice xa13 recessive resistance to bacterial blight is defeated by induction of the disease susceptibility gene Os-11N3. Plant Cell Advance Access published 2010, doi:10.1105/tpc.110.078964.

Aoki, N., Hirose, T., Scofield, G. N., Whitfeld, P. R., and Furbank, R. T. (2003). The sucrose transporter gene family in rice. Plant Cell Physiol. Advance Access published 2003, doi:10.1093/pcp/pcg030.

Bezrutczyk, M., Yang, J., Eom, J. S., Prior, M., Sosso, D., Hartwig, T., Szurek, B., Oliva, R., Vera-Cruz, C., White, F. F., et al. (2018). Sugar flux and signaling in plant–microbe interactions. Plant J. Advance Access published 2018, doi:10.1111/tpj.13775.

Blanvillain-Baufumé, S., Reschke, M., Solé, M., Auguy, F., Doucoure, H., Szurek, B., Meynard, D., Portefaix, M., Cunnac, S., Guiderdoni, E., et al. (2017). Targeted promoter editing for rice resistance to Xanthomonas oryzae pv. oryzae reveals differential activities for SWEET14-inducing TAL effectors. Plant Biotechnol. J. Advance Access published 2017, doi:10.1111/pbi.12613.

Chandran, D. (2015). Co-option of developmentally regulated plant SWEET transporters for pathogen nutrition and abiotic stress tolerance. IUBMB Life Advance Access published 2015, doi:10.1002/iub.1394.

Chen, Q. J., Zhou, H. M., Chen, J., and Wang, X. C. (2006). A Gateway-based platform for multigene plant transformation. Plant Mol. Biol. Advance Access published 2006, doi:10.1007/s11103-006-9065-3.

Chen, R., Zhao, X., Shao, Z., Wei, Z., Wang, Y., Zhu, L., Zhao, J., Sun, M., He, R., and He, G. (2007). Rice UDP-glucose pyrophosphorylase1 is essential for pollen callose deposition and its cosuppression results in a new type of thermosensitive genic male sterility. Plant Cell Advance Access published 2007, doi:10.1105/tpc.106.044123.

Chen, L. Q., Hou, B. H., Lalonde, S., Takanaga, H., Hartung, M. L., Qu, X. Q., Guo, W. J., Kim, J. G., Underwood, W., Chaudhuri, B., et al. (2010). Sugar transporters for intercellular exchange and nutrition of pathogens. Nature Advance Access published 2010, doi:10.1038/nature09606.

Chen, L. Q., Qu, X. Q., Hou, B. H., Sosso, D., Osorio, S., Fernie, A. R., and Frommer, W. B. (2012). Sucrose efflux mediated by SWEET proteins as a key step for phloem transport. Science (80-). Advance Access published 2012, doi:10.1126/science.1213351.

Chen, L. Q., Lin, I. W., Qu, X. Q., Sosso, D., McFarlane, H. E., Londoño, A., Samuels, A. L., and Frommer, W. B. (2015). A cascade of sequentially expressed sucrose transporters in the seed coat and endosperm provides nutrition for the arabidopsis embryo. Plant Cell Advance Access published 2015, doi:10.1105/tpc.114.134585.

Chen, C., He, B., Liu, X., Ma, X., Liu, Y., Yao, H. Y., Zhang, P., Yin, J., Wei, X., Koh, H. J., et al. (2020). Pyrophosphate-fructose 6-phosphate 1-phosphotransferase (PFP1) regulates starch biosynthesis and seed development via heterotetramer formation in rice (Oryza sativa L.). Plant Biotechnol. J. Advance Access published 2020, doi:10.1111/pbi.13173.

Chu, Z., Yuan, M., Yao, J., Ge, X., Yuan, B., Xu, C., Li, X., Fu, B., Li, Z., Bennetzen, J. L., et al. (2006). Promoter mutations of an essential gene for pollen development result in disease resistance in rice. Genes Dev. Advance Access published 2006, doi:10.1101/gad.1416306.

Chung, P., Hsiao, H. H., Chen, H. J., Chang, C. W., and Wang, S. J. (2014). Influence of temperature on the expression of the rice sucrose transporter 4 gene, OsSUT4, in germinating embryos and maturing pollen. Acta Physiol. Plant. Advance Access published 2014, doi:10.1007/s11738-013-1403-x.

Duan, E., Wang, Y., Liu, L., Zhu, J., Zhong, M., Zhang, H., Li, S., Ding, B., Zhang, X., Guo, X., et al. (2016). Pyrophosphate: fructose-6-phosphate 1-phosphotransferase (PFP) regulates carbon metabolism during grain filling in rice. Plant Cell Rep. Advance Access published 2016, doi:10.1007/s00299-016-1964-4.

Eom, J. S., Cho J. Il, Reinders, A., Lee, S. W., Yoo, Y., Tuan, P. Q., Choi, S. B., Bang, G., Park Y. Il, Cho, M. H., et al. (2011). Impaired function of the tonoplast-localized sucrose transporter in rice, OsSUT2, limits the transport of vacuolar reserve sucrose and affects plant growth. Plant Physiol. Advance Access published 2011, doi:10.1104/pp.111.176982.

Eom, J. S., Choi, S. B., Ward, J. M., and Jeon, J. S. (2012). The mechanism of phloem loading in rice (Oryza sativa). Mol. Cells Advance Access published 2012, doi:10.1007/s10059-012-0071-9.

Eom, J. S., Chen, L. Q., Sosso, D., Julius, B. T., Lin, I. W., Qu, X. Q., Braun, D. M., and Frommer, W. B. (2015). SWEETs, transporters for intracellular and intercellular sugar translocation. Curr. Opin. Plant Biol. Advance Access published 2015, doi:10.1016/j.pbi.2015.04.005.

Eom, J. S., Nguyen, C. D., Lee, D. W., Lee, S. K., and Jeon, J. S. (2016). Genetic complementation analysis of rice sucrose transporter genes in Arabidopsis SUC2 mutant atsuc2. J. Plant Biol. Advance Access published 2016, doi:10.1007/s12374-016-0015-6.

Eom, J. S., Luo, D., Atienza-Grande, G., Yang, J., Ji, C., Thi Luu, V., Huguet-Tapia, J. C., Char, S. N., Liu, B., Nguyen, H., et al. (2019). Diagnostic kit for rice blight resistance. Nat. Biotechnol. Advance Access published 2019, doi:10.1038/s41587-019-0268-y.

Feng, B., Sun, Y., Ai, H., Liu, X., Yang, J., Liu, L., Gao, F., Xu, G., and Sun, S. (2018). Overexpression of sucrose transporter OsSUT1 affects rice morphology and physiology. Chinese J. Rice Sci. Advance Access published 2018, doi:10.16819/j.1001-7216.2018.8015.

Gao, Y., Zhang, C., Han, X., Wang, Z. Y., Ma, L., Yuan, D. P., Wu, J. N., Zhu, X. F., Liu, J. M., Li, D. P., et al. (2018). Inhibition of OsSWEET11 function in mesophyll cells improves resistance of rice to sheath blight disease. Mol. Plant Pathol. Advance Access published 2018, doi:10.1111/mpp.12689.

Hirose, T., Takano, M., and Terao, T. (2002). Cell wall invertase in developing rice caryopsis: Molecular cloning of OsCIN1 and analysis of its expression in relation to its role in grain filling. Plant Cell Physiol. Advance Access published 2002, doi:10.1093/pcp/pcf055.

Hirose, T., Zhang, Z., Miyao, A., Hirochika, H., Ohsugi, R., and Terao, T. (2010). Disruption of a gene for rice sucrose transporter, OsSUT1, impairs pollen function but pollen maturation is unaffected. J. Exp. Bot. Advance Access published 2010, doi:10.1093/jxb/erq175.

James, D., Borphukan, B., Fartyal, D., Ram, B., Singh, J., Manna, M., Sheri, V., Panditi, V., Yadav, R., Achary, V. M. M., et al. (2018). Concurrent overexpression of OsGS1;1 and OsGS2 genes in transgenic rice (Oryza sativa L.): Impact on tolerance to abiotic stresses. Front. Plant Sci. Advance Access published 2018, doi:10.3389/fpls.2018.00786.

Jeena, G. S., Kumar, S., and Shukla, R. K. (2019). Structure, evolution and diverse physiological roles of SWEET sugar transporters in plants. Plant Mol. Biol. Advance Access published 2019, doi:10.1007/s11103-019-00872-4.

Jha, G., and Sonti, R. V. (2009). Attack and defense in xanthomonas-rice interactions. Proc. Indian Natl. Sci. Acad. Advance Access published 2009.

Kandoi, D., Mohanty, S. Govindjee, and Tripathy, B. C. (2016). Towards efficient photosynthesis: overexpression of Zea mays phosphoenolpyruvate carboxylase in Arabidopsis thaliana. Photosynth. Res. 130:47–72.

Ke, Y., Hui, S., and Yuan, M. (2017). Xanthomonas oryzae pv. oryzae Inoculation and Growth Rate on Rice by Leaf Clipping Method. BIO-PROTOCOL Advance Access published 2017, doi:10.21769/bioprotoc.2568.

Kim, P., Xue, C. Y., Song, H. D., Gao, Y., Feng, L., Li, Y., and Xuan, Y. H. (2020). Tissue7specific activation of DOF11 promotes rice resistance to sheath blight disease and increases grain weight via activation of SWEET14. Plant Biotechnol. J. Advance Access published 2020, doi:10.1111/pbi.13489.

Kleczkowski, L. A. (1994). Glucose activation and metabolism through UDP-glucose pyrophosphorylase in plants. Phytochemistry Advance Access published 1994, doi:10.1016/S0031-9422(00)89568-0.

Lee, M. W., and Yang, Y. (2006). Transient expression assay by agroinfiltration of leaves. Methods Mol. Biol. Advance Access published 2006, doi:10.1385/1-59745-003-0:225.

Lee, S. K., Jeon, J. S., Börnke, F., Voll, L., Cho J. Il, Goh, C. H., Jeong, S. W., Park Y. Il, Kim, S. J., Choi, S. B., et al. (2008). Loss of cytosolic fructose-1,6-bisphosphatase limits photosynthetic sucrose synthesis and causes severe growth retardations in rice (Oryza sativa). Plant, Cell Environ. Advance Access published 2008, doi:10.1111/j.1365-3040.2008.01890.x.

Lee, S. K., Eom, J. S., Voll, L. M., Prasch, C. M., Park, Y. Il, Hahn, T. R., Ha, S. H., An, G., and Jeon, J. S. (2014). Analysis of a triose phosphate/phosphate translocator-deficient mutant reveals a limited capacity for starch synthesis in rice leaves. Mol. Plant Advance Access published 2014, doi:10.1093/mp/ssu082.

Lemonnier, P., Gaillard, C., Veillet, F., Verbeke, J., Lemoine, R., Coutos-Thévenot, P., and La Camera, S. (2014). Expression of Arabidopsis sugar transport protein STP13 differentially affects glucose transport activity and basal resistance to Botrytis cinerea. Plant Mol. Biol. Advance Access published 2014, doi:10.1007/s11103-014-0198-5.

Li, T., Liu, B., Spalding, M. H., Weeks, D. P., and Yang, B. (2012). High-efficiency TALEN-based gene editing produces disease-resistant rice. Nat. Biotechnol. aAdvance Access published 2012, doi:10.1038/nbt.2199.

Long, W., Dong, B., Wang, Y., Pan, P., Wang, Y., Liu, L., Chen, X., Liu, X., Liu, S., Tian, Y., et al. (2017). FLOURY ENDOSPERM8, encoding the UDP-glucose pyrophosphorylase 1, affects the synthesis and structure of starch in rice endosperm. J. Plant Biol. Advance Access published 2017, doi:10.1007/s12374-017-0066-3.

Ludewig, F., and Flügge, U. I. (2013). Role of metabolite transporters in source-sink carbon allocation. Front. Plant Sci. Advance Access published 2013, doi:10.3389/fpls.2013.00231.

Ma, L., Zhang, D., Miao, Q., Yang, J., Xuan, Y., and Hu, Y. (2017). Essential role of sugar transporter OsSWEET11 during the early stage of rice grain filling. Plant Cell Physiol. Advance Access published 2017, doi:10.1093/pcp/pcx040.

Miao, H., Sun, P., Liu, Q., Miao, Y., Liu, J., Zhang, K., Hu, W., Zhang, J., Wang, J., Wang, Z., et al. (2017). Genome-wide analyses of SWEET family proteins reveal involvement in fruit development and abiotic/biotic stress responses in banana. Sci. Rep. Advance Access published 2017, doi:10.1038/s41598-017-03872-w.

Niittylä, T., Messerli, G., Trevisan, M., Chen, J., Smith, A. M., and Zeeman, S. C. (2004). A Previously Unknown Maltose Transporter Essential for Starch Degradation in Leaves. Science (80-.). Advance Access published 2004, doi:10.1126/science.1091811.

Oliva, R., Ji, C., Atienza-Grande, G., Huguet-Tapia, J. C., Perez-Quintero, A., Li, T., Eom, J. S., Li, C., Nguyen, H., Liu, B., et al. (2019). Broad-spectrum resistance to bacterial blight in rice using genome editing. Nat. Biotechnol. Advance Access published 2019, doi:10.1038/s41587-019-0267-z.

Reidel, E. J., Turgeon, R., and Cheng, L. (2008). A maltose transporter from apple is expressed in source and sink tissues and complements the Arabidopsis maltose export-defective mutant. Plant Cell Physiol. Advance Access published 2008, doi:10.1093/pcp/pcn134.

Ryoo, N., Eom, J. S., Kim, H. B., Vo, B. T., Lee, S. W., Hahn, T. R., and Jeon, J. S. (2013). Expression and functional analysis of rice plastidic maltose transporter, OsMEX1. J. Korean Soc. Appl. Biol. Chem. Advance Access published 2013, doi:10.1007/s13765-012-3266-z.

Sanan-Mishra, N., Pham, X. H., Sopory, S. K., and Tuteja, N. (2005). Pea DNA helicase 45 overexpression in tobacco confers high salinity tolerance without affecting yield. Proc. Natl. Acad. Sci. U. S. A. Advance Access published 2005, doi:10.1073/pnas.0406485102.

Scofield, G. N., Hirose, T., Gaudron, J. A., Upadhyaya, N. M., Ohsugi, R., and Furbank, R. T. (2002). Antisense suppression of the rice sucrose transporter gene, OsSUT1, leads to impaired grain filling and germination but does not affect photosynthesis. Funct. Plant Biol. Advance Access published 2002, doi:10.1071/PP01204.

Scofield, G. N., Hirose, T., Aoki, N., and Furbank, R. T. (2007a). Involvement of the sucrose transporter, OsSUT1, in the long-distance pathway for assimilate transport in rice. J. Exp. Bot. Advance Access published 2007, doi:10.1093/jxb/erm153.

Scofield, G. N., Aoki, N., Hirose, T., Takano, M., Jenkins, C. L. D., and Furbank, R.T. (2007b). The role of the sucrose transporter, OsSUT1, in germination and early seedling growth and development of rice plants. J. Exp. Bot. Advance Access published 2007, doi:10.1093/jxb/erl217.

Singh, J., Pandey, P., James, D., Chandrasekhar, K., Achary, V. M. M., Kaul, T., Tripathy, B. C., and Reddy, M. K. (2014). Enhancing C3 photosynthesis: An outlook on feasible interventions for crop improvement. Plant Biotechnol. J. Advance Access published 2014, doi:10.1111/pbi.12246.

Slewinski, T. L., Meeley, R., and Braun, D. M. (2009). Sucrose transporter1 functions in phloem loading in maize leaves. J. Exp. Bot. Advance Access published 2009, doi:10.1093/jxb/ern335.

Smith, A. M., and Zeeman, S. C. (2006). Quantification of starch in plant tissues. Nat. Protoc. Advance Access published 2006, doi:10.1038/nprot.2006.232.

Sosso, D., Luo, D., Li, Q. B., Sasse, J., Yang, J., Gendrot, G., Suzuki, M., Koch, K. E., McCarty, D. R., Chourey, P. S., et al. (2015). Seed filling in domesticated maize and rice depends on SWEET-mediated hexose transport. Nat. Genet. Advance Access published 2015, doi:10.1038/ng.3422.

Sun, Y., Reinders, A., Lafleur, K. R., Mori, T., and Ward, J. M. (2010). Transport activity of rice sucrose transporters OsSUT1 and OsSUT5. Plant Cell Physiol. Advance Access published 2010, doi:10.1093/pcp/pcp172.

Tanna, B., Choudhary, B., and Mishra, A. (2018). Metabolite profiling, antioxidant, scavenging and anti-proliferative activities of selected tropical green seaweeds reveal the nutraceutical potential of Caulerpa spp. Algal Res. Advance Access published 2018, doi:10.1016/j.algal.2018.10.019.

Tariq, R., Ji, Z., Wang, C., Tang, Y., Zou, L., Sun, H., Chen, G., and Zhao, K. (2019). RNA-Seq analysis of gene expression changes triggered by Xanthomonas oryzae pv. oryzae in a susceptible rice genotype. Rice Advance Access published 2019, doi:10.1186/s12284-019-0301-2.

Wang, L., Lu, Q., Wen, X., and Lu, C. (2015). Enhanced sucrose loading improves rice yield by increasing grain size. Plant Physiol. Advance Access published 2015, doi:10.1104/pp.15.01170.

Woo, M. O., Ham, T. H., Ji, H. S., Choi, M. S., Jiang, W., Chu, S. H., Piao, R., Chin, J. H., Kim, J. A., Park, B. S., et al. (2008). Inactivation of the UGPase1 gene causes genic male sterility and endosperm chalkiness in rice (Oryza sativa L.). Plant J. Advance Access published 2008, doi:10.1111/j.1365-313X.2008.03405.x.

Wu, X., Liu, J., Li, D., and Liu, C. M. (2016). Rice caryopsis development II: Dynamic changes in the endosperm. J. Integr. Plant Biol. Advance Access published 2016, doi:10.1111/jipb.12488.

Wu, Y., Lee, S. K., Yoo, Y., Wei, J., Kwon, S. Y., Lee, S. W., Jeon, J. S., and An, G. (2018). Rice Transcription Factor OsDOF11 Modulates Sugar Transport by Promoting Expression of Sucrose Transporter and SWEET Genes. Mol. Plant Advance Access published 2018, doi:10.1016/j.molp.2018.04.002.

Xu, Z., Xu, X., Gong, Q., Li, Z., Li, Y., Wang, S., Yang, Y., Ma, W., Liu, L., Zhu, B., et al. (2019). Engineering Broad-Spectrum Bacterial Blight Resistance by Simultaneously Disrupting Variable TALE-Binding Elements of Multiple Susceptibility Genes in Rice. Mol. Plant Advance Access published 2019, doi:10.1016/j.molp.2019.08.006.

Yamada, K., Saijo, Y., Nakagami, H., and Takano, Y. (2016). Regulation of sugar transporter activity for antibacterial defense in Arabidopsis. Science (80-.). Advance Access published 2016, doi:10.1126/science.aah5692.

Yang, B., Sugio, A., and White, F. F. (2006). Os8N3 is a host disease-susceptibility gene for bacterial blight of rice. Proc. Natl. Acad. Sci. U. S. A. Advance Access published 2006, doi:10.1073/pnas.0604088103.

Yang, J., Luo, D., Yang, B., Frommer, W. B., and Eom, J. S. (2018). SWEET11 and 15 as key players in seed filling in rice. New Phytol. Advance Access published 2018, doi:10.1111/nph.15004.

Yin, C.-C., Ma, B., Wang, W., Xiong, Q., Zhao, H., Chen, S.-Y., and Zhang, J.-S. (2016). RNA Extraction and Preparation in Rice (Oryza sativa). Curr. Protoc. Plant Biol. Advance Access published 2016, doi:10.1002/cppb.20023.

Yuan, M., and Wang, S. (2013). Rice MtN3/saliva/SWEET family genes and their homologs in cellular organisms. Mol. Plant Advance Access published 2013, doi:10.1093/mp/sst035.

Zaka, A., Grande, G., Coronejo, T., Quibod, I. L., Chen, C. W., Chang, S. J., Szurek, B., Arif, M., Cruz, C. V., and Oliva, R. (2018). Natural variations in the promoter of OsSWEET13 and OsSWEET14 expand the range of resistance against Xanthomonas oryzae pv. Oryzae. PLoS One Advance Access published 2018, doi:10.1371/journal.pone.0203711.

Zeeman, S. C., Smith, S. M., and Smith, A. M. (2004). The breakdown of starch in leaves. New Phytol. Advance Access published 2004, doi:10.1111/j.1469-8137.2004.01101.x.

Zeng, X., Luo, Y., Vu, N. T. Q., Shen, S., Xia, K., and Zhang, M. (2020). CRISPR/Cas9-mediated mutation of OsSWEET14 in rice cv. Zhonghua11 confers resistance to Xanthomonas oryzae pv. oryzae without yield penalty. BMC Plant Biol. Advance Access published 2020, doi:10.1186/s12870-020-02524-y.

