## Supplemental Information for "Coordinated overexpression of *OsSUT1, OsSWEET11* and *OsSWEET14* in rice impairs carbohydrate metabolism that has implications in plant growth, yield and susceptibility to *Xanthomonas oryzae pv oryzae (Xoo)*"

### Supplemental Data

**Supplemental Figure 1. (A and B)** PCR amplification of *OsSWEET11* and *OsSWEET14* nucleotide fragments used in the study **(C)** PCR amplification of 1617 bp *OsSUT1* CDS from rice cDNA.; **(D)** PCR amplification of ~1.6Kb promoter region of OsSUT1; **(E)** Steps of multiround gateway technology to stack all the three gene cassettes in plant transformation vector *pMDC99*; **(F)** MCS of EV-1 and 2.

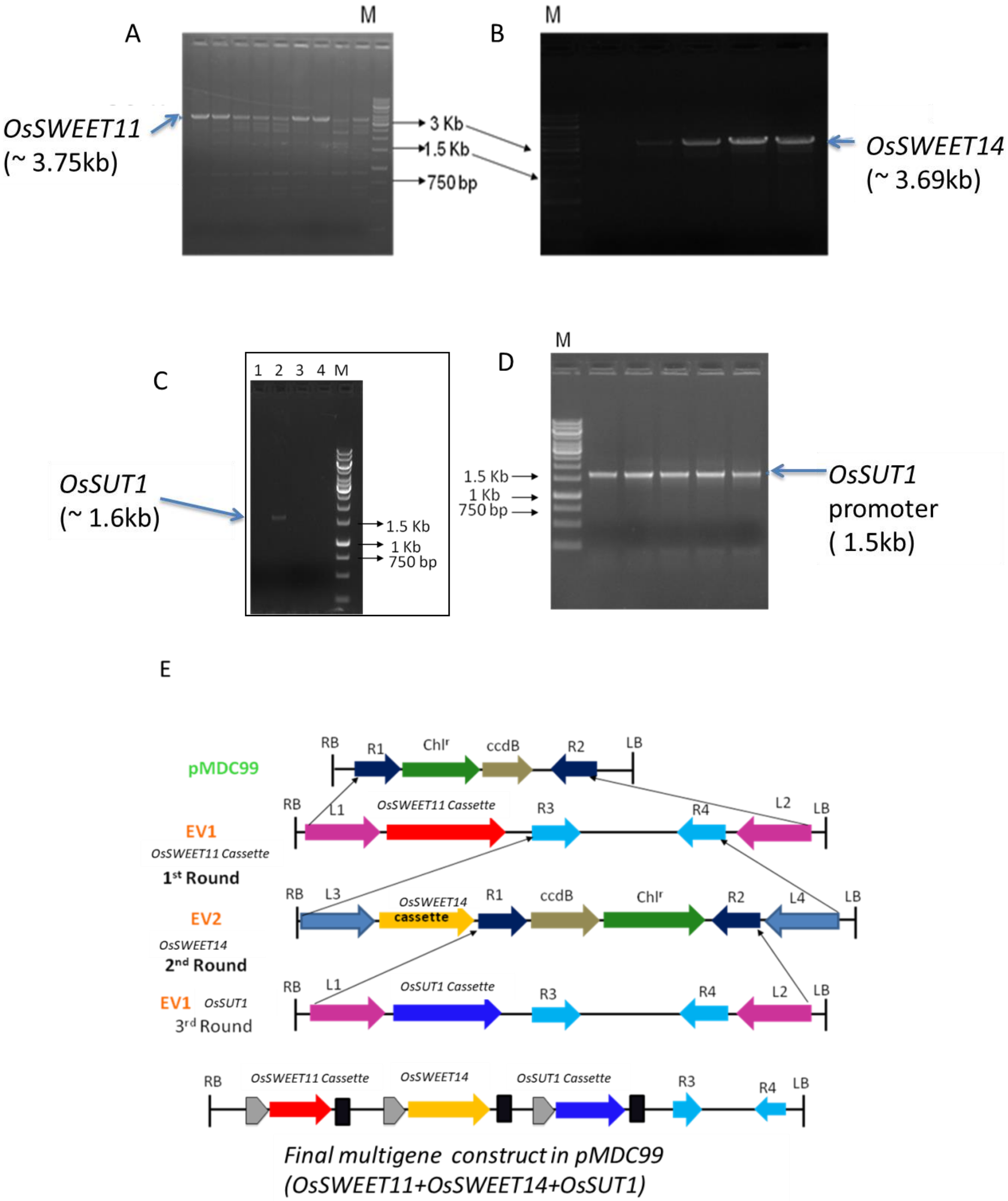

F

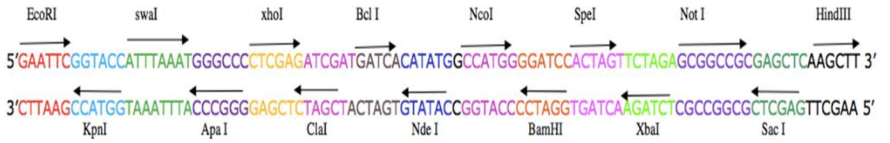


**Supplemental Figure 2. (A)** Genomic DNA PCR for screening of putative $T_0$ transgenic plants harboring the recombinant construct with all the three gene cassettes; **(B)** Southern hybridization of different PCR positive transgenic plants with *HPTII* as a probe; **(C)** RT-PCR of transgenic lines SW5, SW9 and SW8.

A

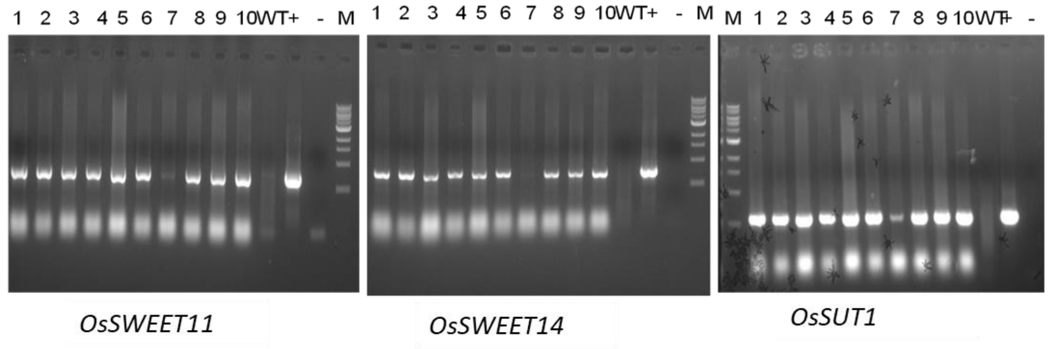


B

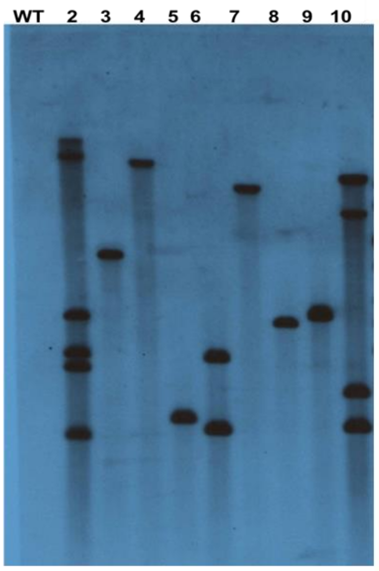


C

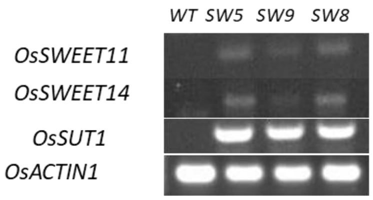

**Supplemental Figure 3. (A)** GUS visualization in tobacco leaves. –ve control is leaf from WT tobacco plants, test is transgenic tobacco leaf transformed with transcriptional fusion construct *pOsSWEET11:GUS*, +ve control is *GUS* under *CaMV 35S* promoter; **(B)** Enlarged view of test leaf; **(C)** Transient expression of *pOsSWEET14:GUS* in tobacco leaf with +ve control; **(D)** Enlarged view of *pOsSWEET14:GUS* tobacco leaf.

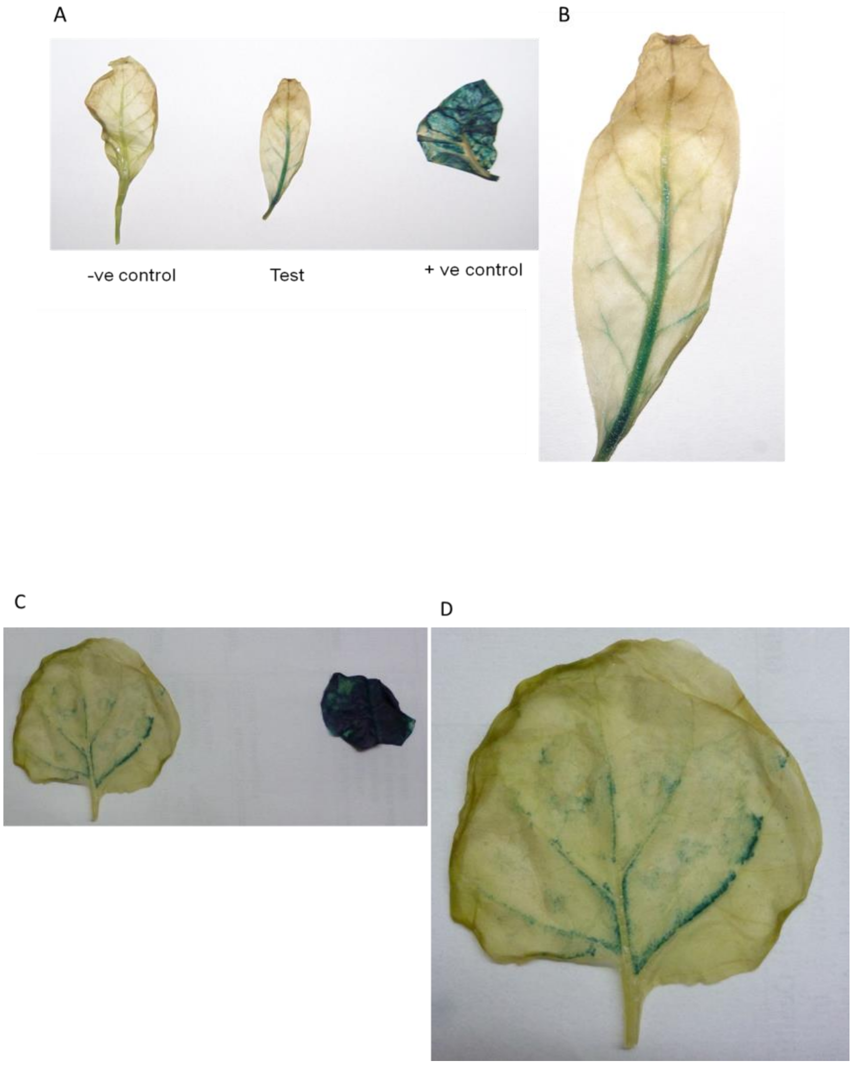

### Supplemental Table 1

List of primers and Gene ids:

| Gene Name | Locus id | Primer sequence Forward 5'-3' | Reverse 5'-3' |
|---|---|---|---|
| *OsSWEET11* | LOC_Os08g42350 | AA**GGATCC**CGATCAGGTACCAAATCTTT | TA**GCGGCCGC**TCACACGGCGGCGGTGATCT |
| *OsSWEET14* | LOC_Os11g31190 | AA**GGATCC**CTTTCTAGTTCATATTAATT | TA**GCGGCCGC**TCAGTGACCGCCGCCCATGC |
| *OsSWEET11* for promoter cloning in GUS vector | | AA**GGATCC**CGATCAGGTACCAAATCTTT | TTGTTAAC TGCTACTGGTGATGAAGGTTA |
| *OsSWEET14* for promoter cloning in GUS vector | | AA**GGATCC**CTTTCTAGTTCATATTAATT | ATGTTAAC TGCAGCAAGATCTTGATTAAC |
| *OsSUT1* for Promoter cloning | | TT**CTCGAG**TAAAATGTGAGGCCGACATG | AA**GGATCC**GGCGGACGCGCCACGCACAA |
| *OsSUT1* for CDS | LOC4331775/LOC_Os03g07480.2 | AA**GGATCC**ATGGCTCGCGGCAGCGGGGC | TA**GCGGCCGC**TCAGTGACCGCCGCCCATGC |
| *CaMV 35S* Terminator | | TA**GCGGCCGC**TCGCTGAAATCACCAGTCTC | TT**GTCGAC**GATCTGGATTTTAGTACTGG |
| Primers for gDNA screening of transgenic plants with gene forward and *CaMV 35S* terminator reverse | | | |
| *OsSWEET11* | | ATCAAGACCAAGAGCGTCGAGTTCAT | CGAAACCCTATAAGAACCCTAATT |
| *OsSWEET14* | | AACTAAGGATGCACATATTTACGCA | CGAAACCCTATAAGAACCCTAATT |
| *OsSUT1* | | AGCTCATTCTTGATTGAACCAATG | CGAAACCCTATAAGAACCCTAATT |
| *HPTII* | | ATGAAAAAGCCTGAACTCACC | CTATTTCTTTGCCCTCGGAC |
| For RT-PCR | | | |
| *OsSWEET11* | | TGGTTCTGCTACGGCCTCTT | AGAGAGAGACTGGTGATTTC |
| *OsSWEET14* | | AATATGTCGCTCTTCCCAACGT | AGAGAGAGACTGGTGATTTC |
| *OsSUT1* | | AGGGAACTGATCCTCAGATCGA | AGAGAGAGACTGGTGATTTC |
| *OsACTIN1* | | GCCGTCCTCTCTCTGTATGC | GCAATGCCAGGGAACATAGT |
| Primers for RT-qPCR | | | |

| | | | |
|---|---|---|---|
| *OsSWEET11* | | TGGTTCTGCTACGGCC TCTT | GGTACCAGAAGTAG AGCCCCATCT |
| *OsSWEET14* | | ATCAAGCCTTCAAGCA AAGC | CTAGGAGACCAAAG GCGAAG |
| *OsSUT1* | | TGGCCAAGGGCTCTGC A | GATGAGCGCGAATC CCGAG |
| *OsSWEET4* | LOC_Os02g19820 | CGCTCGCATGATCGTCG GTAT *TCCACGCTCTTAGTCTGGATCAC* | TCCACGCTCTTAGTC TGGATCAC |
| *OsSWEET15* | LOC_Os02g30910 | CCGTACGTGGTGACGC TCTT | ACGCACCCCACACCA TTG |
| *OsSWEET13* | LOC_Os12g29220 | CTACGCGCTGATCAAG TCCAA | GGGCGTAGGCGAGG TACAT |
| *OsSWEET6b* | LOC_Os01g42090 | GGAAGGACGTGGAGC AGTTC | GCTGTTCGGGTGGAC GAT |
| *OsACTIN1* | LOC_Os03g50885 | CCCCCATGCTATCCTT CGT | GGCCGTTGTGGTGAA TGAGT |
| *cyOsFBP1* | LOC_Os01g64660 .1 | CGCACCTGCGTTCTTG TCT | AGCCGTCGAGTGGAT CAAAG |
| *OsGlcT* | LOC_Os01g04190 .1 | ACGTGGAGCACTTGGT TCTGT | CCACCAAGCAGGGTT TCCT |
| *OsGPT1* | LOC_Os08g08840 | TGCACAGAGCGTTTTC TACCA | ACGCTTCATCGTATT GCCAAT |
| *OsMT1* | LOC_Os04g51330 | CTTGCAGTGCTTCCTC AGGTT | CAGCTGTGGCAGCCA AAGA |
| *OsPFP* | LOC_Os02g48360 | GCATTCGAGCTATTGA GGAACA | TTGAATCAGCACCTG CTCCTT |
| *OsPPase* | LOC_Os01g64670 | CTCCGGCTGTGTTCAA CGTT | AAGACCCGATCAAC CATGATG |
| *OsSPS1* | LOC_Os01g50730 / Os01g0702900 | AGTTCGTGGCGATAAC ATGGA | GCCAGGCATCATTGC AAGT |
| *OsTPT1* | LOC_Os01g13770 | AAGTTCCCCTGCCTCT TTGG | TTGATGAAGCCCGTC CAGTT |
| *OsTPT2* | LOC_Os05g15160 | CCGTGTCCTTTGCTGCT GTA | TGCTGGCCAAGGATA AACTGT |
| *OsSS1* | LOC_Os06g09450 | TGGTGACCACGGCAAT CA | GCGGATATGCCCCTT CAACT |
| *OsSS2* | LOC_Os03g28330 | GAGCTGGATTTCGAGC CATTC | TCGATGACAGATGCC TGTTGA |
| *OsSS3* | LOC_Os07g42490 | AACTTTCTTCGTGCGC ACAAC | TTTCTGCCTTCCTCA ATGCA |

| | | | |
|---|---|---|---|
| *OsUGPase1* | LOC_Os09g38030 | GTGATGTGTTCCCCTCCTTGA | GCACCCAAGTTGTCCGAGTT |
| *OsSUT2* | LOC_Os12g44380 | AACCGGCATTGTCATTGCTT | AACCCGACTAGCAGCCATTG |
| *OsSUT3* | LOC_Os10g26470 | CGACTGAGGACTGCAAGGTTT | TGCACGGTGTTGTTGGAGAA |
| *OsSUT4* | LOC_Os02g58080 | ACAACGGTGTCCGAGAAGGT | GCACCCATCAGTCGGCATA |
| *OsSUT5* | LOC_Os02g36700 | CCAGGCCGTGTTGTTCAGT | GGCAATGTTGAGGACACCAAT |
